# Negative Binomial factor regression with application to microbiome data analysis

**DOI:** 10.1101/2021.11.29.470304

**Authors:** Aditya K. Mishra, Christian L. Müller

**Author notes:** **Correspondence** Aditya K. Mishra, Center for Computational Mathematics, Flatiron Institute, Simons Foundation, New York, NY 10010.

## Abstract

The human microbiome provides essential physiological functions and helps maintain host homeostasis via the formation of intricate ecological host-microbiome relationships. While it is well established that the lifestyle of the host, dietary preferences, demographic background, and health status can influence microbial community composition and dynamics, robust generalizable associations between specific host-associated factors and specific microbial taxa have remained largely elusive. Here, we propose factor regression models that allow the estimation of structured parsimonious associations between host-related features and amplicon-derived microbial taxa. To account for the overdispersed nature of the amplicon sequencing count data, we propose Negative Binomial reduced rank regression (NB-RRR) and Negative Binomial co-sparse factor regression (NB-FAR). While NB-RRR encodes the underlying dependency among the microbial abundances as outcomes and the host-associated features as predictors through a rank-constrained coefficient matrix, NB-FAR uses a sparse singular value decomposition of the coefficient matrix. The latter approach avoids the notoriously difficult joint parameter estimation by extracting sparse unit-rank components of the coefficient matrix sequentially. To solve the non-convex optimization problems associated with these factor regression models, we present a novel iterative block-wise majorization procedure. Extensive simulation studies and an application to the microbial abundance data from the American Gut Project demonstrate the efficacy of the proposed procedure. In the American Gut Project data, we identify key factors that strongly link dietary habits and host life style to specific microbial families.

## 1 INTRODUCTION

The human microbiome, the collection of microbes that reside on or within human tissues and fluids, has formed intricate ecological relationships with the host over the course of co-evolution^1^. Advances in next-generation amplicon sequencing technology and analysis techniques have enabled the direct identification of microbial species compositions and abundances in their natural habitat. These approaches have revealed considerable variability in both composition and diversity across different body sites ^2^ and allowed the estimation of potential associations between the microbiome and the underlying health condition of the subject^3^. For instance, differential abundances of the gut microbiome have been linked to medical conditions such as inflammatory bowel disease (IBD), irritable bowel syndrome (IBS), Type 2 diabetes, obesity, and neurological disorders^4^. The gut microbiome also makes considerable contribution to metabolic functioning, e.g., by fermenting non-digestible substrates such as fibers and endogenous mucus^5^. Dietary changes can thus induce considerable shifts in gut microbial compositions^6^. Similar intricate relationships between the environment and the microbiome are known in other ecosystems as well. For instance, the soil microbiome plays a significant role in the cycle of carbon and nitrogen fixation, thus having direct implications for plant growth^7^. However, soil microbiome compositions show large variability with respect to soil conditions such as pH, temperature, moisture, and spatial location. Likewise, cyanobacteria in the marine ecosystems contribute to a large extent to the ocean’s primary productivity, yet show considerable abundance variability across location, season, and water conditions^8,9^.

Amplicon-based microbiome survey data are derived from samples of the habitat of interest, e.g., the human gut, where variable regions of the bacterial and archaeal 16S ribosomal RNA are experimentally extracted and sequenced. These marker gene sequences serve as a proxy to the underlying bacterial taxon abundances and are summarized in Operational Taxonomic Units (OTUs) or Amplicon Sequence Variants (ASVs)^10,11^. Reference databases are used to identify the (approximate) taxonomy of the representative microbial sequences. Standard bioinformatic workflows and databases, such as, e.g., QIIME-2^10^ or the Qiita framework^12^, allow standardized processing of and access to these OTU/species counts. In addition, large-scale microbiome survey studies such as the American Gut Project (AGP)^13^, the Human Microbiome Project (HMP)^14^, and the Earth Microbiome Project ^15^ also collect host- or environment-associated covariate data that can be large/high-dimensional. These survey data reach a level of scale and completeness that, in principle, allows to make quantitative predictions about the relationship between host-associated factors and microbial abundance patterns. For example, the AGP data comprises hundreds of host-associated features, including variables indicating dietary intake, medical conditions, medication use, participants’ demography, and life style.

Using the AGP data as representative microbiome data resource, we here introduce a statistical factor regression framework that allows the identification of key associations between host-related features and microbial taxa. While recent work^16^ has already identified individual host factors that confound microbial abundance patterns in relation to specific disease phenotypes, we propose a general factor model that simultaneously takes into all relevant host-associated covariates and links them to the observed microbial abundances, independent of a specific downstream task. Since the observed microbial abundances across all levels of taxonomic aggregation come in form of overdispersed count data, we base our model on the classical Negative Binomial (NB) distribution (see Figure 1). Negative Binomial models are common place in genomics and microbiome data analysis. For example, the popular DESeq2 package^17^, used extensively in differential expression testing in bulk RNA-Seq data, uses the Negative Binomial model as underlying model for total mRNA transcript abundance. Due to technical and experimental limitations, microbial count data, however, carry only relative information and show varied sequencing depth across samples. To mitigate these limitations, microbiome data require transformation/normalization approaches prior to statistical modeling^18^. Important examples include rarefying samples to a common sequencing depth or scaling using factors such as a cumulative sum, median, upper quartile, or the total sum, the latter of which leading to compositional or relative abundance data. A particularly popular approach for Negative Binomial modeling of microbial count data is common sum scaling, as put forward by McMurdie and Holmes (2014)^19^. When modeling microbial count data on the OTU/ASV level, zero-inflated extensions of the NB model have been proposed to account for the excess number of zeros in the data^20^ (see, e.g.,^21^ for a critical assessment).

**FIGURE 1.**
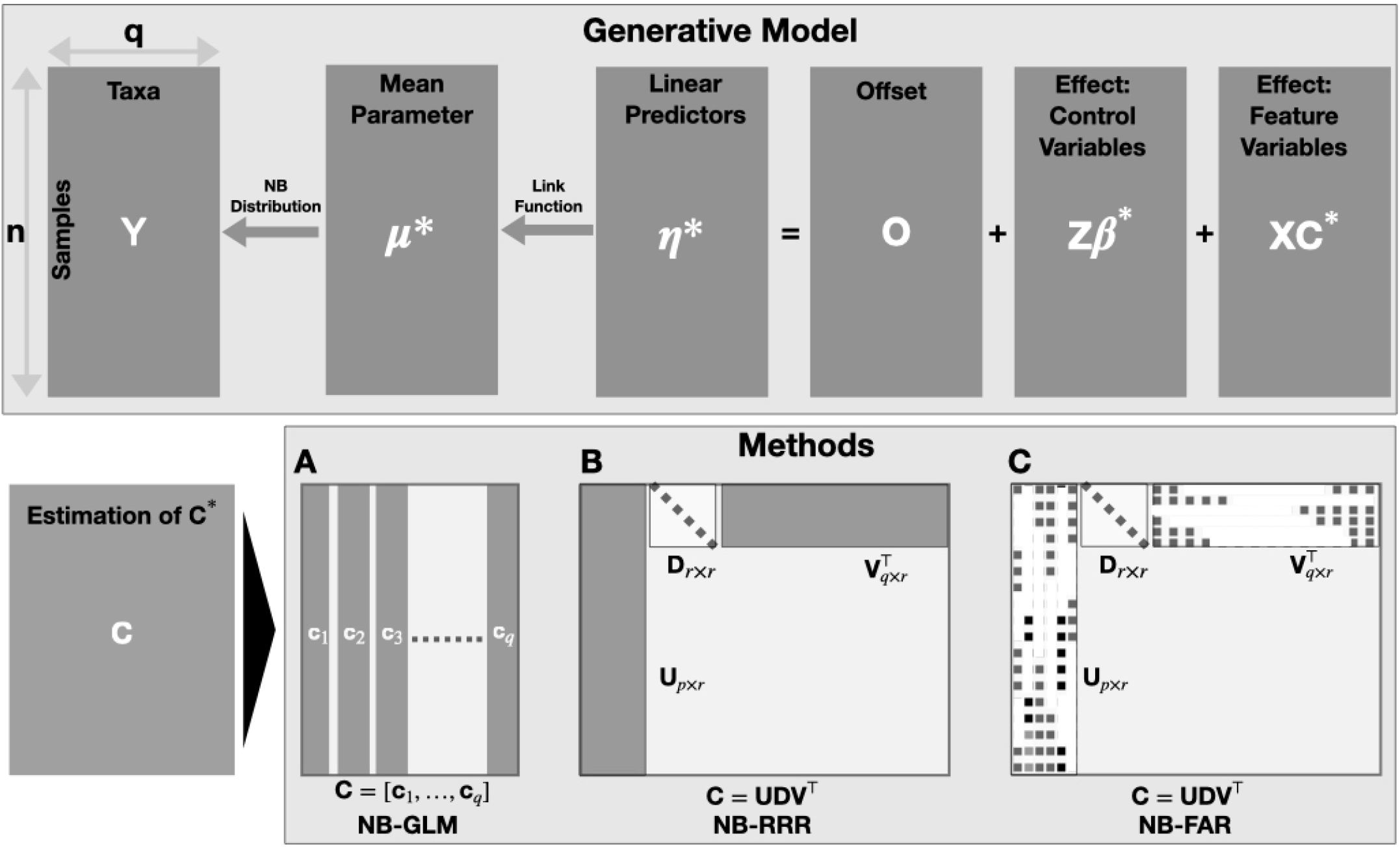
Regression model for the overdispersed microbial abundance data **Y** of count types in terms of the covariates **X** and the control variables **Z**. The upper panel presents the true generative model with parameters {***β***\*, **C**\*} and the lower panel presents the three approaches for estimating the parameters. A) The Generalized linear model of Negative Binomial regression (NB-GLM) estimates each of the columns of **C** = [**c**_1_, …, **c**_*q*_] separately; B) Negative Binomial reduced rank regression (NB-RRR) jointly estimates a low-rank coefficient matrix as **C** = **UDV**^T^; C)Negative Binomial co-sparse factor regression (NB-FAR) jointly estimates a low-rank coefficient matrix as **C** = **UDV**^T^ with sparse singular vectors {**U,V**}.

Alternative generative statistical modeling approaches include the Dirichlet Multinomial (mixture) framework^22^, latent Dirichlet allocation ^23,24^, and Poisson distribution models^25^ (including their respective zero-inflated extensions). Several models also allow the inclusion of host or environmental covariate data in generative modeling, including Poisson factor models^26,27,28^, latent Dirichlet allocation ^29^, and Bayesian Dirichlet multinomial models^30,31^. The majority of these procedures, however, tackles the arguably more challenging problem of modeling data at the amplicon or species level. Since the majority of the OTUs/ASVs are absent in most of the samples, the addition of zero-inflated components complicates model estimation, and the dimensionality of the data makes biological interpretability challenging.

Our modeling framework proposes a different strategy, namely, the accurate identification and estimation of key host-associated factors that influence microbial taxa at higher taxonomic ranks. This approach allows crisp, biologically interpretable statements about the underlying potential role of host features on broad microbial abundance patterns at the expense of sacrificing taxonomic resolution. However, even when summarizing microbiome survey data on a higher taxonomic level, the data remains overdispersed but does not contain excess zeros. This makes the addition of zero-inflated components in the model superfluous (see Figure S5 of the supplementary materials). Prior work^32^ already established that Negative Binomial regression (NB-GLM) is capable of relating individual taxa to the host-associated features (see Figure 1(A)). This marginal model, however, ignores the fact that some of the taxa are likely influenced by a set of common factors, e.g., age, diet, or life style.

Here, we alleviate this shortcoming and introduce a Negative Binomial factor regression framework that models microbial abundance data as outcome and covariates as predictors jointly while encoding the underlying dependencies in a parsimonious fashion. In the high-dimensional multivariate linear regression setting, it is common practice to model the underlying dependency for dependent outcomes via a structured coefficient matrix ^33^ (see Figure 1(B,C)). The model parameters are estimated by solving a regularized optimization problem. For example, reduced-rank regression^34,35,36^ promotes information-sharing among response and predictors through a low-rank coefficient matrix. When covariates are high-dimensional, sparsity is known to facilitate identifiability and better model interpretation^37^. In the multivariate setting, this has been achieved via a sparse factorization of the model coefficient matrix ^38,39,40,33^. When the outcome matrix comprises non-Gaussian or mixed type variables, e.g., Bernoulli-type for binary outcomes and Poisson-type for counts, Luo et al. (2017)^41^ proposed *mixed-outcome reduced-rank regression*. Mishra et al. (2020)^42^ proposed *generalized co-sparse factor regression* (GO-FAR) to model the outcome jointly under sparsity constraints. As we will show in the reminder of the manuscript, these existing models are inappropriate for microbial taxon data due to the overdispersed nature of the counts.

Instead, we propose *Negative Binomial reduced rank regression* (NB-RRR) and *Negative Binomial co-sparse factor regression* (NB-FAR) to jointly model the microbial abundance data using the host-associated features as covariates (see Figure 1(B-C) for details). NB-RRR follows previous reduced rank regression frameworks by capturing the underlying dependencies among response and predictors via a low-rank coefficient matrix. NB-FAR extends the GO-FAR framework and encodes the underlying dependency via a sparse singular value decomposition (SSVD) of the coefficient matrix. Following the estimation strategy of GO-FAR, we extract unit-rank components of the coefficient matrix sequentially^33^, thus alleviating the challenging problem of joint estimation. There, each sequential step solves a co-sparse unit rank estimation problem with a suitably adjusted offset term that accounts for the effects of previous steps. NB-FAR thus models the associations of microbial abundance and host-associated features via a few latent factors that comprise only a subset of predictors. Both NB-RRR and NB-FAR estimation procedures are implemented, tested, validated, and made publicly available in the R package `nbfar`, available on GitHub at [LINK].

The remainder of the paper is organized as follows. Section 2 provides the details of the NB-RRR and NB-FAR framework. Section 3 provides the details of the parameter estimation procedure. In Section 4, we present simulation studies to demonstrate the efficacy of the estimation procedures. Section 5 provides a detailed analysis of the AGP taxon data on the family level and an extensive set of host-associated covariates using our NB factor regression methods. Section 6 discusses the findings and provides future research directions. Additional data analysis plots and the details of the estimation procedures are provided in the Supplementary Materials.

## 2 FACTOR MODELS FOR MICROBIOME DATA

As motivating example we consider the data from the American Gut Project (AGP)^13^ where samples from thousands of participants have been collected, sequenced, and processed to obtain microbial abundances. Each sample is associated with participant-specific covariates that are related to diet, heath, and lifestyle. Our overall goal is to understand the associations of these covariates with the observed microbial abundance patterns. Let us denote the abundance/count data of *q* taxa from *n* samples as **Y** = [*y*_*ik*_]_*n*×*q*_ = [**y**_1_, … **y**_*n*_]^T^ ∈ ℝ^*n*×*q*^, the associated predictors/covariates as **X** = [**x**_1_, … **x**_*n*_]^T^ ∈ ℝ^*n*×*p*^, and the control variables as **Z** = [**z**_1_, … **z**_*n*_]^T^ ∈ ℝ^*n*×*c*^. **Z** comprises variables, such as age and gender, that are held constant in an experiment and are thus fully adjusted for in the model.

The observed taxon abundance data are overdispersed, i.e., the variance of the taxa tends to be considerably larger than their mean^43^. This fact motivates the use of a parametric framework based on the Negative Binomial distribution to model the underlying associations between multivariate count outcome **Y** and factors {**X**, **Z**}. Using the alternative parametrization of the Negative Binomial distribution^44^, the generative model for the abundance of the *j*th taxon in the *i*th sample is given by

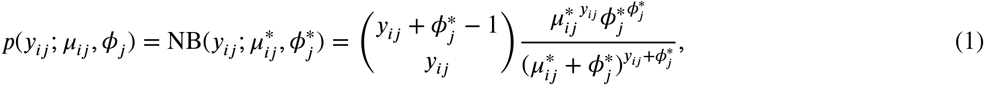

where 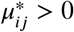 is the entry-specific mean parameter and 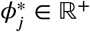 is the taxon-specific shape parameter. Let us jointly represent the shape parameters of *q* taxa by 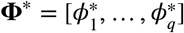 and the entry-specific mean parameters by 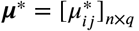. For the generative model (1), 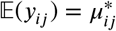 and 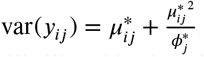, i.e., var(*y*_*ij*_) ≥ 𝔼(*y*_*ij*_), making the model suitable for overdispersed count data. Then, the joint negative log-likelihood is given by

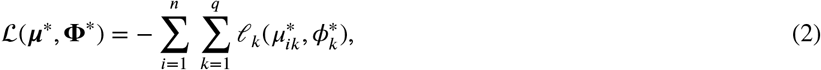

where 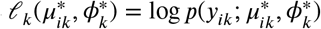.

To associate the participant-specific covariates to the microbial abundance, we link entry-specific mean parameters ***μ***\* to the linear predictors as

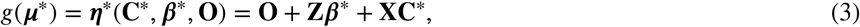

where *g*(·) is a suitable link function that satisfies ***μ***\* *>* 0, **O** = [*o*_*ik*_]_*n×q*_ ∈ ℝ^*n*×*q*^ is a fixed offset term, 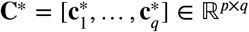 is the coefficient matrix corresponding to the predictors **X**, and 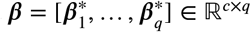 is the coefficient matrix corresponding to the control variables **Z**. The formulation includes an intercept in the model by setting the first column of **Z** to be **1**_*n*_, the *n* × 1 vector of ones. Following the work of Zeileis et al. (2008)^44^ and Anders et al. (2010)^45^, we choose *g*(*x*) = log *x* as the link function so that any 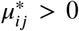. Depending on the problem, one may choose another link function that satisfies ***μ***\* *>* 0. Unless otherwise stated, we write ***η***\* (**C**\*, ***β***\*, **O**) as ***η***\*.

With the shape parameter fixed, the Negative Binomial distribution (1) belongs to the exponential dispersion family^46^. To utilize the form of this family, we define the corresponding natural parameter matrix 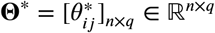 as

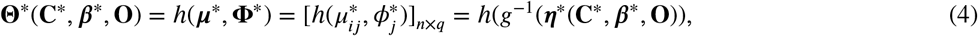

where 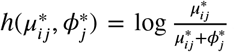. Again, unless otherwise stated, we will conveniently express **Θ**\* (**C**\*, ***β***\*, **O**) as **Θ**\*. We assume the outcomes to be conditionally independent given **X** and **Z**. In terms of the natural parameter **Θ\***, we rewrite the negative log-likelihood function (2) as

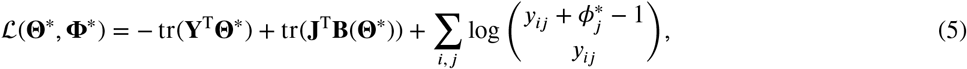

where **J** =**1**_*n*×*q*,_ tr(**A**) is the *trace* of a square matrix **A** and 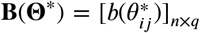 such that 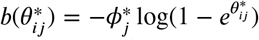. For fixed shape parameter **Φ\***, it is straightfoorward to show that 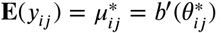. This results in linking **Θ\*** to the linear predictor ***η***\* via 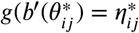.

To obtain an estimate of the model parameters, we minimize the objective function **ℒ**(**Θ,Φ**) with respect to {**C**,***β***,**Φ**}, where *g*(*b*′(**Θ**(**C, *β***,**O**))) = ***η***(**C**,***β***,**O**) = **O** + **XC**+**Z*β*** and **Φ** = [*ϕ*_1_, …, *ϕ*_*j*_]. Since it is likely that taxa in *Y* may be biologically related due to factors such as a co. Also, the data problem may have large/high dimensional predictors. Hence, the multivariate model has correlated predictors and interrelated responses. Minimizing **ℒ**(**Θ,Φ**) with respect to {**C**,***β***,**Φ**} is equivalent to separately fitting a Negative Binomial regression model for each outcome, which ignores the dependency among covariates and responses. Building upon the recent advances in the multivariate mixed outcomes modeling approach taken by Luo et al. (2017)^41^ and Mishra et al. (2020)^42^, we encode the dependency using low-rank and SSVD of the coefficient matrix.

The microbial abundance data model (1) with the rank constraint

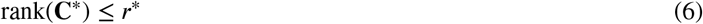

is referred to as **N**egative **B**inomial **R**educed **R**ank **R**egression, denoted **NB-RRR**. The low-rank coefficient matrix **C**\* implies a significantly lower number of effective parameters that exhibit better estimation performance in the high dimensional data problems^33,42,38^. Any *low-rank* coefficient matrix **C**\* can be expressed as the product of any two low-rank matrices. Based on the formulation of the linear predictor ***η***\* (3), we associate the responses to latent factors that are constructed as linear combinations of predictors **X**. However, in the large dimensional setting, only a subset of predictors are relevant, and the latent factors may be associated with only a subset of responses. This can be achieved by expressing **C**\* as a product of two unique and identifiable low-rank matrices that are entrywise sparse. Motivated by the recent work of Mishra et al. (2017)^33^ and Mishra et al. (2020)^42^, we assume that the singular value decomposition (SVD) of the coefficient matrix **C**\* in (1) is *co-sparse*^33^ and decompose **C**\* as

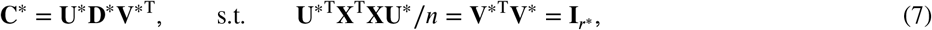

where both the left singular vector matrix 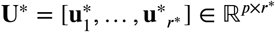 and the right singular vector matrix 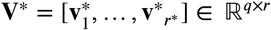 are assumed to be *sparse*, and 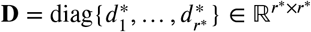 is the diagonal matrix with the nonzero singular values on its diagonal. In the high-dimensional setting, this formulation facilitates better interpretation. Additional orthogonality constraints in the formulation ensure that the SVD of **C**\* is identifiable and the latent factors 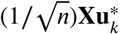 for *k* = 1 …***r***\* are uncorrelated. Each of the right singular vectors in **V**\* associates the corresponding latent factor to the microbial abundance outcome **Y**, and the singular values 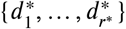 denote the strength of the association. We thus term the proposed model **N**egative **B**inomial *co-sparse* **Fa**ctor **R**egression, denoted by **NB-FAR**.

Since the majority of the available microbial count data only provides relative abundance information, it is common practice to scale or transform the data prior to statistical analysis ^19^. Finding a suitable normalization/transformation approach remains an active area of research in microbiome data analysis ^19,18,47^. NB-RRR and NB-FAR enable proper scaling of the data by specifying an offset **O** ∈ ℝ^*n*×*q*^ matrix with *o*_*ik*_ denoting the *ik*th entry. Throughout this manuscript, we use common sum scaling^19,18^ as the default setting to scale the microbial count data **Y** to the minimum sequencing depth. This corresponds to setting *o*_*ik*_ = 0 in our model. However, inspired by the differential abundance analysis tool DESeq2^17^, NBFAR’s implementation enables the user to choose other offset terms or to specify them manually using prior knowledge. Specifically, one can use

a. *o*_*ik*_ = 0 when the observed abundance data is normalized using common sum scaling^18^,
b. 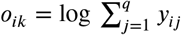 for total sum scaling,
c. 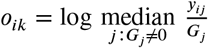 where 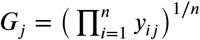 for median scaling,
d. 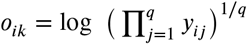 for centered-log-ratio scaling.

Each of the normalization approaches via the offset term will, however, influence the outcome of the analysis ^18^.

## 3 ESTIMATION PROCEDURES

Parameter estimation in both models requires minimizing a constrained, non-convex negative log-likelihood function **ℒ**(**Θ,Φ**) with respect to {**C**,***β***,**Φ**}. Joint estimation of the parameters {**C**,***β***,**Φ**} satisfying the rank constraint (6) in the case of NB-RRR and the orthogonality constraints (7) in the case of NB-FAR is a notoriously difficult problem. In both cases, we solve the optimization problem using the majorization-minimization (MM) approach^48^, an alternating procedure that updates the blocks of parameters in cyclic order until convergence. In an update step, we minimize a convex surrogate that majorizes the objective function of the optimization problem.

### 3.1 Negative Binomial Reduced Rank Regression (NB-RRR)

The optimization problem to estimate the parameters of NB-RRR is given by

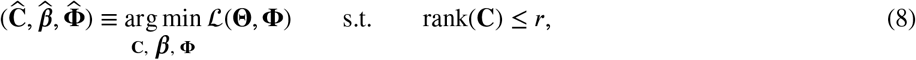

where *g*(*b*′(**Θ**(**C**,***β***,**O**))) = ***η***(**C**,***β***,**O**) = **O** + **XC**+**Z*β***. Let us denote the problem as NB-RRR(**C**,***β***,**Φ**; **W, Z, X, O**,***r***). Unless otherwise stated, we will write **Θ(C**,***β*, O**) as **Θ** and ***η***(**C**,***β*, O**) as ***η***. Using the framework of MM, we minimize the objective function using an iterative procedure that cycles between **C**-step, ***β***-step, and **Φ**-step to update **C, *β*** and **Φ**, respectively, until convergence.

In the **C**-step, for fixed ***β*** and **Φ**, let us denote the natural parameter **Θ** and **ℒ** (**Θ**,**Φ**) as **Θ**(**C**) and **ℒ**(**Θ**(**C**)), respectively. Suppose differentiable **ℒ**(**Θ**(**C**)) is L-Lipschitz continuous gradient function for some constant *L*_*c*_ i.e., ∥ **▽ ℒ** (**Θ**(**Č**)) − **▽ ℒ**(**Θ**(**C**)) ∥ ≤ *L*_*c*_∥**Č** −**C**∥ for any conformable **Č**. The statement holds for any *L*_*c*_ such that sup_**C**_ ∥ **▽**^2^**ℒ**(**Θ**(**C**))∥≤ *L*_*c*_ = max_1≤*j≤q*_∥**X**^T^diag(**Y**_.***j***_+ 1)**X**∥/2; see Section 1.1 of the Supplementary Material for details. Using the result, we majorize **ℒ**(**Θ**(**C**), **Φ**) by a convex surrogate at a given **Č** and update the parameter **C** as 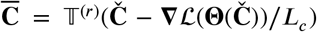, where 𝕋^(*r*)^(**M**) extracts *r* SVD components of matrix **M**. Similarly, in the ***β***-step, for fixed **C** and **Φ**, we denote **ℒ**(**Θ**,**Φ**) as **ℒ**(**Θ**(***β***)). Following the **C**-step procedure, **ℒ**(**Θ**(***β***)) is also L-Lipschitz continuous gradient function for some constant *L*_*b*_ = max_1≤*j*≤*q*_∥**Z**^T^diag(**Y**_.***j***_+ 1)**Z**∥/2; see Section 1.1 of the Supplementary Material for details. At any 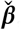, we majorize **ℒ**(**Θ**(***β***)) and then update the parameter ***β*** as 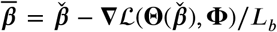. Finally, in the **Φ**-step, we follow Zeileis et al. (2008)^44^ to update each shape parameter *ϕ*_*j*_ using the Newton-Raphson method for fixed **C** and ***β***; see Section 1.6 of the Supplementary Material for details. We compute the parameter estimates for 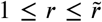 where 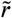 is the user-specified conservative maximum rank, and select a rank *r* using K-fold cross-validation ^49^. The iterative procedure is summarized in Algorithm 1.

#### 3.1.1 Monotonically decreasing property of NB-RRR

Using Algorithm 1, we estimate the parameters {**C**,***β***,**Φ**,} of NB-RRR. The iterative procedure consists of minimizing several convex surrogates of the objective function with fixed Lipschitz constants {*L*_*b*_, *L*_*c*_}. Let us jointly denote the updated parameters from **C**-step, ***β***-step and **Φ**-step after *t*th iteration by {**C**^(*t*)^, ***β***^(*t*)^, **Φ**^(*t*)^}.

##### Theorem 1

The sequence of parameter estimates {**C**^(*t*)^,***β***^(*t*)^,**Φ**^(*t*)^}_*t*∈ℕ_ obtained using Algorithm 1 satisfies

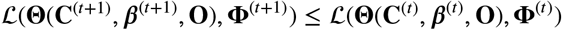

for the constants *L*_*b*_ = max_1≤*j*≤*q*_∥**Z**^T^diag(**Y**_.***j***_+ 1)**Z**∥/2 and *L*_*c*_ = max_1≤*j*≤*q*_∥**X**^T^diag(**Y**_.***j***_+ 1)**X**∥/2.

We have relegated the proof of Theorem 1 to Section 1.2 of the Supplementary Material. In extensive simulation studies, we have found that the sequence always converges in practice.

##### Algorithm 1

Negative Binomial Reduced Rank Regression (NB-RRR)

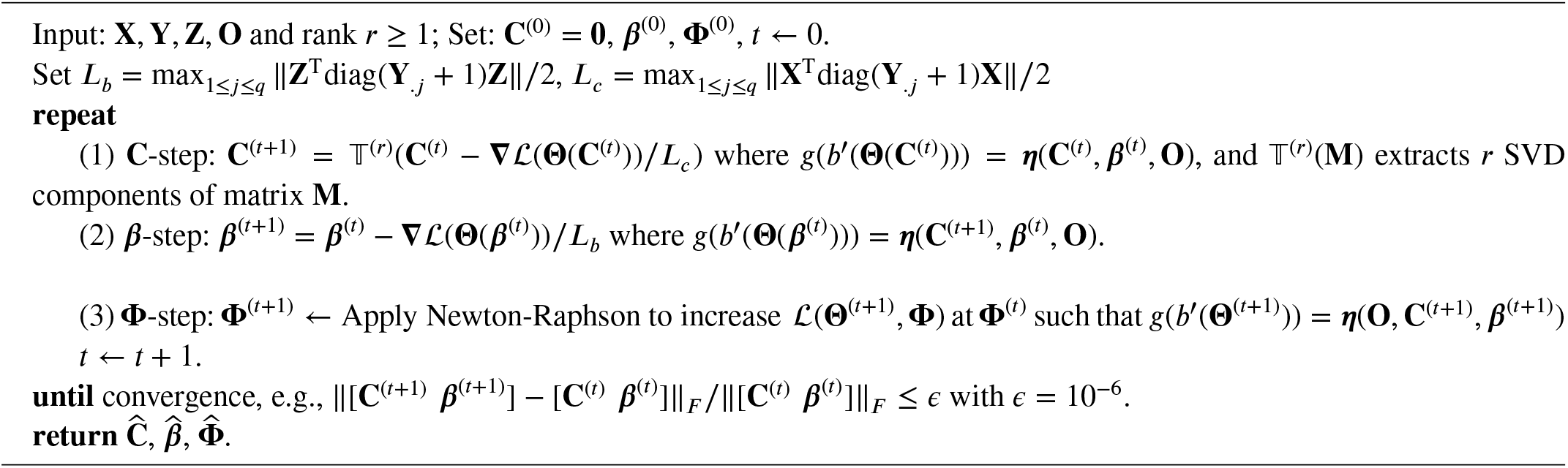

### 3.2 Negative Binomial Co-Sparse Factor Regression (NB-FAR)

Joint estimation of the model parameters {**U, D, V**,***β***,**Φ**} requires solving an optimization problem that minimizes **ℒ**(**Θ**,**Φ**) in the presence of a sparsity-inducing penalty on {**U, V**} and the orthogonality constraint (7) ^42^. The optimization problem requires the rank *r* to be specified. Since existing optimization tools are computationally inefficient for the task, we extend the sequential extraction procedure, proposed by Mishra et al. (2017,2020)^33,42^, for NB-FAR and estimate the unit-rank components of 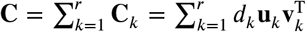, i.e., (*d*_*k*_,**u**_*k*_,**v**_*k*_), for *k* = 1, …, ***r***. Let **Ĉ**_*i*_ for *i* = 1, …, *k*− 1 be the estimate of the unit-rank components. Then, to extract the *k*th unit-rank component in the *k*th step of the sequential procedure, we solve the optimization problem

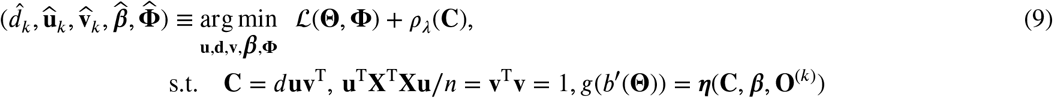

where *ρ*_*λ*_(**C**) is a sparsity-inducing penalty function with tuning parameter *λ* and 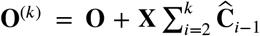 with **O**^(1)^ = **O** is the offset term. This problem is referred to as **N**egative **B**inomial **C**o-sparse **U**nit-**R**ank **E**stimation (NB-CURE) with input parameters **C**,***β***,**Φ;Y**,**X**,**Z**,**O**^(*k*)^ and penalty function *ρ*, in short, NB-CURE (**C**,***β***,**Φ;Y**,**X**,**Z**,**O**^(*k*)^, *ρ*).

Following Mishra et al. (2020)^42^, we use the elastic net penalty and its adaptive version^50^ for the *k*th step as

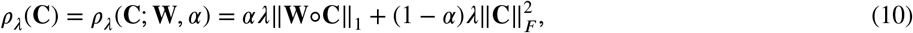

where the operator “ ○ ” stands for the Hadamard product, **W** = [*w*_*ij*_]_*p*×*q*_ is a pre-specified weighting matrix, *λ* is a tuning parameter controlling the overall amount of regularization and *α* ∈ (0, 1) controls the relative weights between the two penalty terms. In the *k*th step of NB-FAR, we let 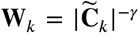, where *γ* = 1 and 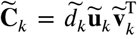 is an initial estimate of **C**_***k***_. Here, we solve NB-RRR **(C**,***β***,**Φ;Y**,**Z**,**X**,**O**^(*k*)^,1) to obtain this initial estimate 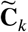 and extract 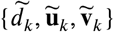. Assuming that the NB-CURE problem can be solved for a suitable tuning parameter *λ* (see Sec. 3.2.1), NB-FAR’s estimation procedure is summarized in Algorithm 2.

#### 3.2.1 Computation of Negative Binomial Constrained Unit-Rank Regression (NB-CURE)

The general form of the optimization problem for the *k*th step of the sequential procedure, i.e., NB-CURE (**C**,***β***,**Φ;Y**,**Z**,**X**,**O**,*ρ*), is given by

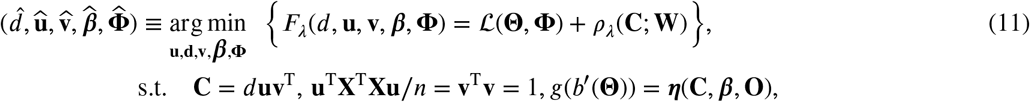

where **W** = *w*^(*d*)^**w**^(*u*)^**w**^(*υ*)T^. In practice, we fix *α* = 0.95 and denote *ρ*_*λ*_(**C**; **W**, *α*) as *ρ*_*λ*_(**C**; **W**). Similar to NB-RRR model estimation, we use the MM framework and solve the optimization problem using an iterative procedure that cycles between **u**-step, **v**-step, ***β***-step and **Φ**-step to update the parameters in blocks of (**u**,*d*), (**v**,*d*), ***β*** and **Φ**, respectively, until convergence.

##### Algorithm 2

Negative Binomial Co-Sparse Factor Regression (NB-FAR)

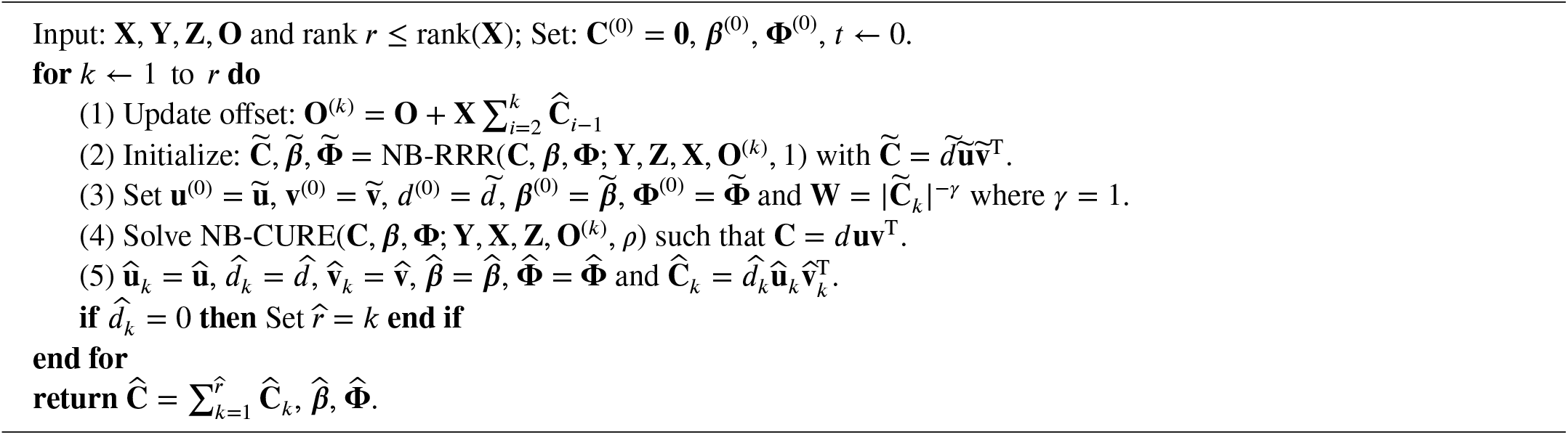

Let us represent **Θ** and 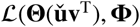, which are functions of **C**, as **Θ**(**C**) and 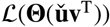, respectively. In the **u**-step, for fixed 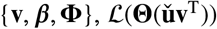 has L-Lipschitz continuous gradients for some ***L***_*u*_ where 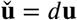. Again, using this fact and following the NB-RRR estimation procedure, we majorize 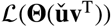 by a convex surrogate and then minimize it to update the block variable 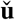 as

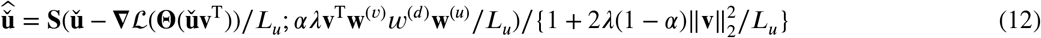

where 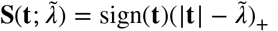 is the element-wise soft-thresholding operator on any **t** ∈ ℝ^*p*^; see Section 1.3 of the Supplementary Material for details. Similarly, in the **v**-step, for fixed {**u**,***β***,**Φ**} with 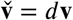, we show that 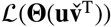 is a L-Lipschitz continuous gradient function for some *L*_*υ*_ and update the block variable 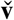 as

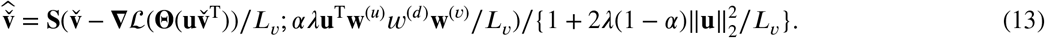

We apply the equality constraints in (9) to recover the estimate of {*d*,**u**,**v**} from the estimate of the block variables 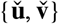. In the ***β***-step and the **Φ**-step, we follow the corresponding step of NB-RRR parameter estimation (see Algorithm 1) to update ***β*** and **Φ**, respectively. We have relegated the details of the convex surrogate function that majorizes the objective function, the computation of the constants (*L*_*u*_, *L*_*υ*_, *L*_*b*_), and the update steps to Section 1.3 of the Supplementary Material.

We compute the parameter estimates for several *λ* values (the default is 50) in the range of *λ*_*max*_ to *λ*_*min*_ that are equi-spaced on a log scale, where *λ*_*max*_ = 2∥**X**^T^(**Y**− *g*^−1^(**O**))∥_∞_ and *λ*_*min*_ = 1*e*^−6^ × *λ*_*max*_. We apply K-fold cross-validation^49^ to select a tuning parameter *λ*. The iterative procedure is summarized in Algorithm 3.

##### Algorithm 3

Negative Binomial Constrained Unit-Rank Regression (NB-CURE)

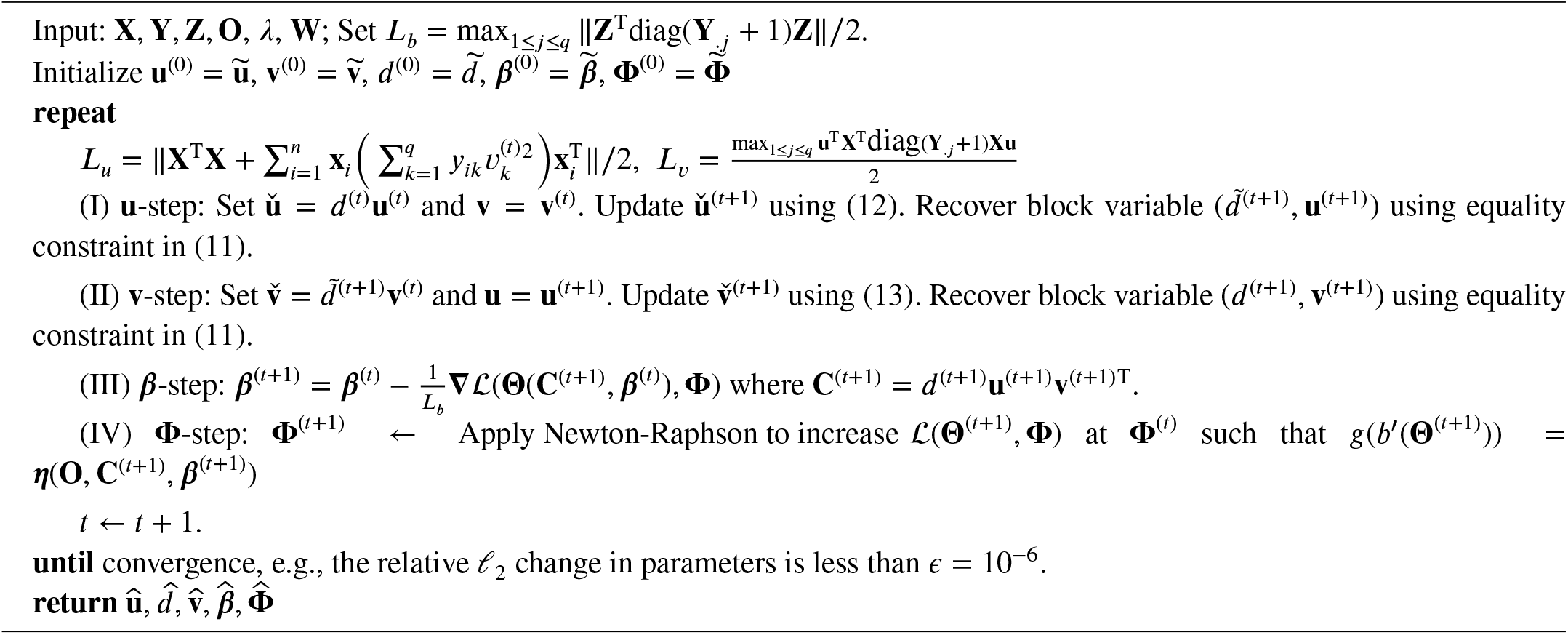

#### 3.2.2 Monotonically decreasing property of NB-CURE

Using Algorithm 3, we solve the optimization problem of NB-CURE to estimate the parameters {**d**,**u**,**v**,***β* Φ**,}. In the Iterative Procedure, the objective function (11) is majorized by a convex surrogate in each of the **u**-step, **v**-step, and ***β***-step, and then minimized. Let us jointly denote the updated parameters from **u**-step, **v**-step, ***β***-step and **Φ**-step after *t*th iteration by {*d*^(*t*)^,**u**^(*t*)^,**v**^(*t*)^,***β***^(*t*)^,**Φ**^(*t*)^}.

##### Theorem 2

The sequence of parameters estimate {*d*^(*t*)^, **u**^(*t*)^,**v**^(*t*)^,***β***^(*t*)^,**Φ**^(*t*)^}_*t*∈ℕ_ obtained from Algorithm 3 satisfies

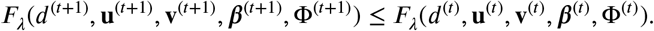

for 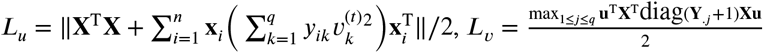 and *L*_*b*_ = max_1≤*j≤q*_∥**Z**^T^diag(**Y**_.***j***_+ 1)**Z**∥/2.

We have relegated the proof of Theorem 2 to Section 1.4 of the Supplementary Material. Similar to NB-RRR, we have found in extensive simulation studies that the sequence always converges in practice.

#### 3.2.3 Handling missing outcome values in NB-FAR

Besides microbiome data, the NB factor models may also prove useful for multivariate count data in other domains, including genomics, sports, image analysis, and text mining. A common scenario in these domains is the presence of missing entries in the outcome matrix **Y**. To highlight NB-FAR’s ability to account for missing entries, we can extend the framework of (5) by calculating the negative log-likelihood as follows.

Let us define an index set of the observed entries in **Y** as **Ω** = {(*i,k*); *y*_*ik*_ is observed,*i* = 1,…*n,k* = 1,…,*q*}, and denote the projection of **Y** onto **Ω** by 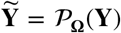, where 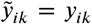 for any (*i,k*) **∈ Ω** and 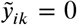 otherwise. Following Mishra et al. (2020)^42^, we write the negative log-likelihood function with incomplete data as

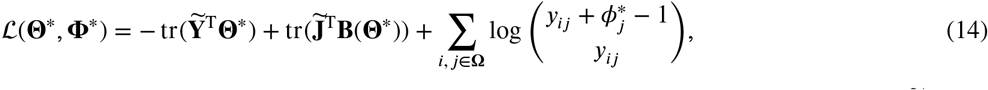

where 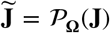 and *g*(*b*′(**Θ**^**\***^) = ***η\****. In case of missing entries in the outcome matrix **Y**, one should replace **Y** with 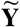 and **J** with 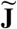 and apply our proposed procedure to estimate the parameters. The same approach is applicable in the NB-RRR model.

## 4 SIMULATION STUDIES

### 4.1 Setup

We compare the performance of NB-RRR and NB-FAR with GO-FAR and NB-GLM to showcase the efficacy of the proposed procedures in modeling multivariate overdispersed count data in the high/large-dimensional settings. We evaluate the performance of the methods in terms of estimation error, prediction accuracy, sparsity recovery, rank identification, and shape error. GO-FAR (implemented in the R package `gofar`) assumes that the underlying distribution of the count outcomes is Poisson. The specific comparison with GO-FAR allows us to probe the effect of overdispersion in the data on model quality. The comparison with NB-GLM highlights the potential limitations of marginal approaches in modeling dependent variables.

We followed Mishra et al. (2020)^42^ to simulate the predictor matrix **X** and sparse SVD components of the coefficient matrix **C\***, i.e., {**U*, D*, V\***}. Our setup considers the true rank ***r***\* = 3 with 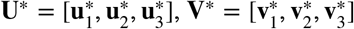 and 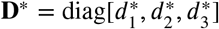 such that 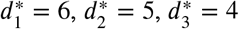. The notations unif(𝒜, *b*) denote a vector of length *b* whose entries are uniformly distributed on the set 𝒜 and rep(*a, b*) denote the vector of length *b* with all entries equal to *a*. We generate 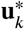 as 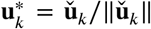, where 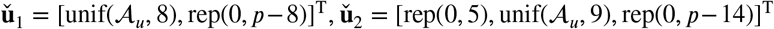, and 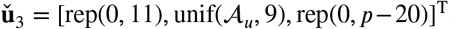. Similarly, we generate 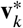 as 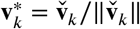, where 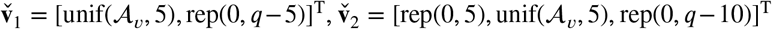 and 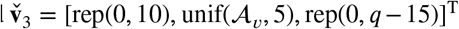. Here we set 𝒜_*u*_= ±1 and 𝒜_*υ*_,= [−1, −0.3]∪[0.3, 1]. An intercept is included in the model by setting **Z** =**1**_*n*_ with ***β***^*^= [rep(0.5,*q*)]^T^. We have considered simulation settings with *p* = 100 and *p* = 300 to demonstrate the efficacy of the proposed procedure in large/high-dimensional examples.

We simulate the predictor matrix **X** ∈ ℝ^*n*×*p*^ from a multivariate normal distribution with some rotations such that the latent factors 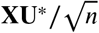 satisfy the orthogonality constraint (7); refer to the simulation study of Mishra et al. (2017)^33^ for details on the formulation. At the OTU/ASV level in the taxonomy, typical microbial abundance observations are excessively sparse. Since our factor models are tailored toward modeling taxa aggregated on a higher taxonomic rank, e.g., the family level, we first estimate the level of sparsity in the observed AGP data. We found that, on the family level, 20% of the entries AGP data are zeros. In the simulation setting, we thus set the shape parameters to 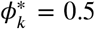 for *k* = 1,…*q*,resulting in 20% zero entries in the simulated outcome matrix **Y**. Based on the model suggested in (1), we simulate **Y** such that *g*(***μ****\**) = ***η\****= **Z*β\**** + **XC\***. Finally, we also include a simulation scenario where 20% of entries in the response matrix **Y** are missing at random. The latter scenario showcases the ability of NB-FAR and NB-RRR to handle missing values in **Y**.

We evaluate model performance in terms of a) the estimation error Er(**C**) = ∥**Ĉ** − **C\***∥_*F*_/(*pq*), b) the prediction error 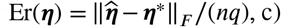 sparsity recovery using the false positive rate (FPR) and the false negative rate (FNR) and d) rank estimation 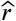 FNR is computed by comparing the support (non-zero entries) of 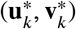 with corresponding entries in its estimate 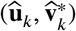 for *k* = 1,…*r\**. The FPR, on the other hand, compares zero entries in 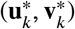 with corresponding entries in its estimate 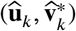 for *k* = 1, …, *r\**. In case of overestimated rank, we report the relative residual signal in the excessive components as 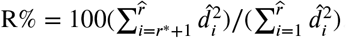. The error in the shape parameter estimate is reported as 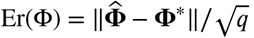.

### 4.2 Results

Since we observed similar model performances in the *p* = 100 and *p* = 300 scenarios, we report the model evaluation statistics only for the latter case in Table 1. The results are average performances over 100 replicates. The model comparison for the *p* = 100 case is available in Table S1 of the Supplementary Material. The boxplot in Figure 2 compares the models in terms of prediction error Er(***η***) (see Figure S1 of the Supplementary Material for comparison on the basis of estimation error Er(**C**)). Compared to the standard approach of modeling overdispersed count outcome using the marginal Negative Binomial regression model (NB-GLM) or the Poisson counterpart of NB-FAR, i.e., GO-FAR, both Negative Binomial Factor models show superior performance in terms of parameter estimation, prediction, support identification, and rank estimation. In the present model scenario, NB-FAR outperforms NB-RRR at the expense of a higher computational cost. Since the true parameters of the underlying model are sparse, we expect and confirm this superior behavior of NB-FAR. The performance decrease of the GO-FAR model highlights the effect of the misspecification in the model with respect to the overdispersed data. In particular, the performance of GO-FAR considerably deteriorates in terms of false negative rate. Based on the results in the simulation examples where 20% of entries in **Y** are missing (M%20 rows in Table 1), we observe that NB-FAR can efficiently estimate the model parameters with slight deterioration compared to the full data model.

**TABLE 1.** Simulation: model evaluation based on 100 replications using various performance measures (standard deviations are shown in parentheses) in case of p = 300 with negative binomial responses.

| | M% | Er(C) | Er($\Theta$) | FPR | FNR | R% | r | Er($\Phi$) | Time(s) |
| --- | --- | --- | --- | --- | --- | --- | --- | --- | --- |
| NB-FAR | 0 | 2.68 (1.10) | 20.70 (3.32) | 5.10 (1.60) | 1.42 (1.44) | 0.00 (0.00) | 3.00 (0.00) | 0.10 (0.07) | 263.20 (28.01) |
| NB-RRR | 0 | 14.49 (1.14) | 40.99 (1.71) | 100.00 (0.00) | 0.00 (0.00) | 0.00 (0.00) | 3.00 (0.00) | 0.63 (0.14) | 95.63 (8.43) |
| GO-FAR | 0 | 10.79 (2.17) | 47.58 (5.77) | 4.56 (3.58) | 49.80 (30.21) | 0.00 (0.00) | 2.51 (0.98) | 1.37 (0.00) | 60.49 (27.45) |
| NB-GLM | 0 | 276.31 (10.27) | 163.75 (2.15) | 100.00 (0.00) | 0.00 (0.00) | 69.66 (1.29) | 29.58 (0.56) | 3741.83 (149.34) | 1376.36 (35.38) |
| NB-FAR | 20 | 3.36 (1.35) | 22.98 (3.58) | 5.63 (2.13) | 2.51 (2.29) | 0.00 (0.00) | 3.00 (0.00) | 0.12 (0.08) | 273.12 (26.96) |
| NB-RRR | 20 | 15.15 (1.35) | 43.77 (2.07) | 100.00 (0.00) | 0.00 (0.00) | 0.00 (0.00) | 3.00 (0.00) | 0.74 (0.17) | 92.31 (8.80) |
| GO-FAR | 20 | 11.78 (2.59) | 50.37 (7.45) | 4.89 (3.79) | 54.17 (28.03) | 0.00 (0.00) | 2.54 (0.99) | 1.37 (0.00) | 79.46 (20.03) |
| NB-GLM | 20 | 228.27 (7.35) | 172.59 (2.53) | 100.00 (0.00) | 0.00 (0.00) | 71.53 (1.24) | 29.68 (0.47) | 3773.07 (135.35) | 1774.59 (41.64) |

**FIGURE 2.**
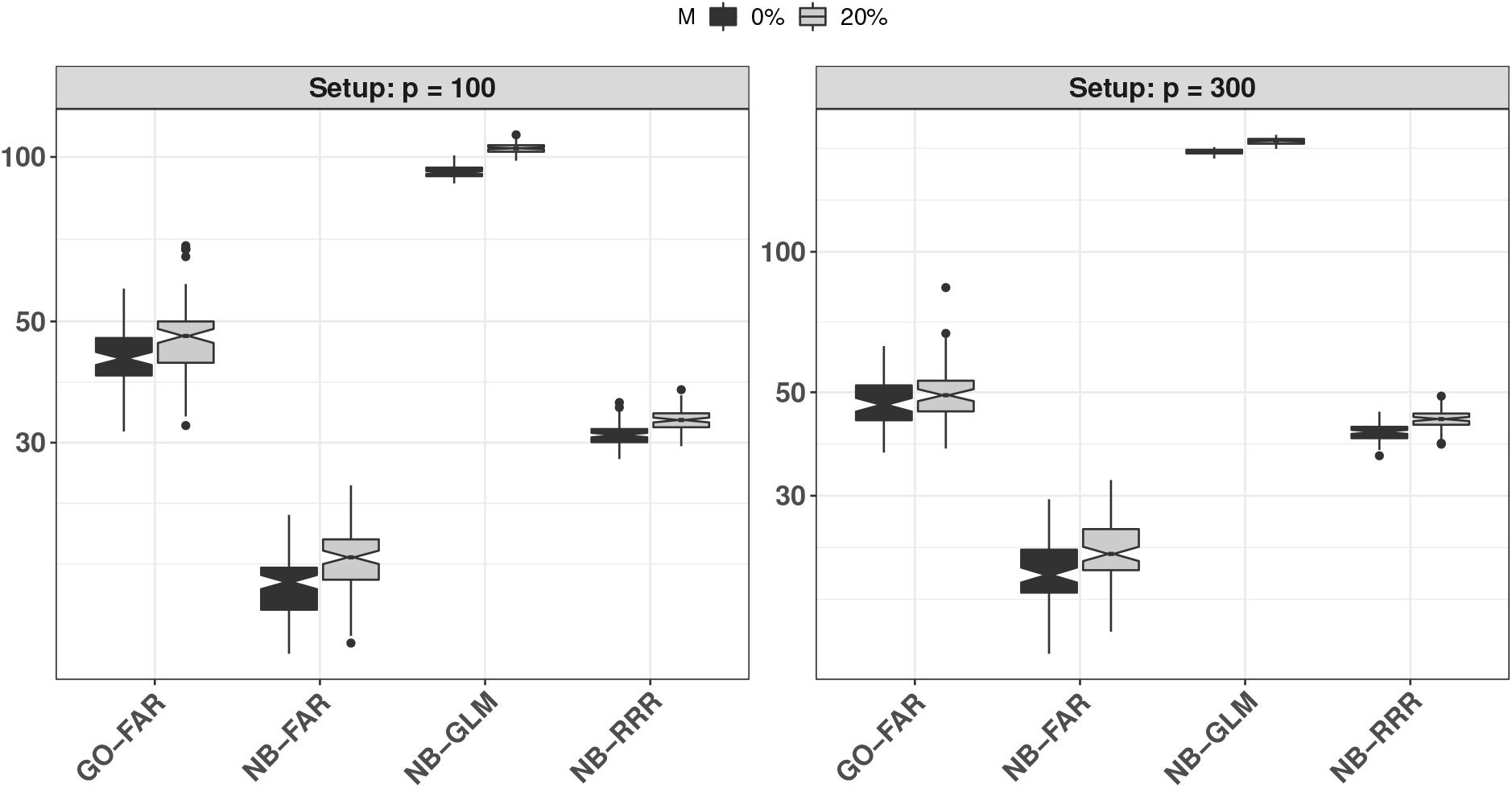
Notched boxplots of the prediction error Er(***η***) on the simulated count data under Setup I (*p* = 100) and II (*p* = 300), respectively. The results are based on 100 replications.

## 5 APPLICATION

We now illustrate the performance of the Negative Binomial factor models using the microbial abundance data from the American Gut Project (AGP)^13^. AGP comprises around 30,000 fecal, oral, hand, skin, and other body site samples. We used Qiita^12^, an open-source platform for microbial study, to access the raw data. We selected and downloaded the microbial abundance data that were obtained after processing 150-nucleotides-long trimmed sequences from the 16S V4 region and used the provided Greengenes reference database for taxonomic annotation. The AGP study also records metadata/covariates related to each participant’s health condition, diet, demography, nutrient intake, and habit. Here, we focused on a subset of *n* = 627 participants where both fecal amplicon data (with sufficient sequencing depth *>* 2000) and VioScreen variables were available. The VioScreen variables provide a detailed account of the dietary habits of the participants.

We aggregated the microbial count data to the family level using the available taxonomic annotation and performed common sum scaling^19,18^ at the minimum sequencing depth. After dropping taxa observed in less than 10% of samples, we arrive at *q* = 39 microbial families as multivariate outcome **Y**. As described in the Methods part, we apply common sum scaling to the outcome matrix Y to arrive at a uniform sampling depth per sample^19^. We curated the metadata as follows. First, we dropped several descriptive variables, such as, e.g., sample name and sample identifier. We then removed all variables that were missing in more than 50 out of *n* = 627 samples. The final covariate matrix **X** comprises *p* = 357 predictors. We manually assigned each of the different predictors to high-level categories, such as, e.g., diet, habit, health-related, nutrient-related, etc. The curated AGP dataset with the assigned categories is available within the R package (see also simulationExamples.R in the Supplementary Material for the sample data obtained after pre-processing). Finally, we considered gender, body mass index (BMI), and age as control variables **Z**. We also included an intercept leading to **Z** ∈ 627 × 4.

We used NB-FAR and NB-RRR to learn about the underlying association between the set of *p* = 357 covariates and the observed microbial family abundances with a special focus on the patterns in the low-rank and sparse coefficient matrix estimates **C**. For comparison, we also ran NB-GLM and GO-FAR and assessed model quality based on AIC, BIC, and log-likelihood. We also report rank estimates *r* and model size. Table 2 summarizes model performance results from 100 replications with 80% of data used for training and 20% for testing. Compared to the marginal approach of NB-GLM and the GO-FAR model, NB-FAR and NB-RRR achieve considerably lower AIC, BIC, and log-likelihood. Both NB-FAR and NB-RRR have comparable prediction errors on the test data and choose approximately rank 3 models. Interestingly, GO-FAR estimates a rank-1 model, though at the expense of decreased predictive performance.

**TABLE 2.** Summary of average model performances on the AGP data in terms of AIC, BIC, and log-likelihood (second to forth column) on the test data. The other columns summarize the average rank estimate ***r*** and support of the singular vectors of **C** as {Supp(**U**), Supp(**V**)}.

| Model | AIC | BIC | Log-likelihood | $r$ | Supp($\mathbf{U}$) | Supp($\mathbf{V}$) |
| --- | --- | --- | --- | --- | --- | --- |
| NB-FAR | 1.37 | 1.71 | -9.44 | 3.15 | 7.97 | 16.80 |
| NB-RRR | 1.86 | 3.69 | -9.23 | 3.15 | 100.00 | 100.00 |
| GO-FAR | 34.90 | 35.20 | -19.00 | 1.00 | 17.90 | 3.59 |
| NB-GLM | 36.40 | 37.10 | -19.80 | 27.20 | 55.10 | 86.00 |

Since NB-FAR achieved comparable performance to NB-RRR in terms of predictive log-likelihood with considerably reduced model complexity, we re-estimated the model parameters of NB-FAR on the full data set using Algorithm 2. NB-FAR again identified a rank(**C**) = 3 solution, enabling a parsimonious and interpretable description of host covariates -microbial family associations with only three sparse latent factors, given by sparse left and right singular vectors **Û** and **V**. The support size (% of non-zero entries) of the estimates of the singular vectors are supp(**U**) = {16%, 17%, 20%} and supp(**V**) = {28%, 74%, 59%}. Using the union of the support of the estimated **U** and **V**, we can visualize all associations with the block-sparse coefficient matrix **C**, shown in Figure 3(left panel). The other panels in Figure 3 display the individual unit-rank components **C**_1_, **C**_2_, and **C**_3_. Note that each of the unit-rank components is orthogonal to one another. We highlight the high-level categories of the covariates, including diet, habit, and health, by vertical lines. Horizontal lines delineate the different microbial phyla to which the families belong to. We observed that each of the singular vectors induced a different sparse pattern of positive (red) and negative (blue) blocks of associations between covariates and taxa. Overall, we found that 30 out of the 39 microbial families were found to be associated with host-associated covariates. The families *Pseudomonadaceae, Barnesiellaceae, Paraprevotellaceae, Christensenellaceae, Coriobacteriaceae, ML615J-28, Mogibacteriaceae, Ys2*, and *Oxalobacteraceae* were not associated with any of the covariates.

**FIGURE 3.**
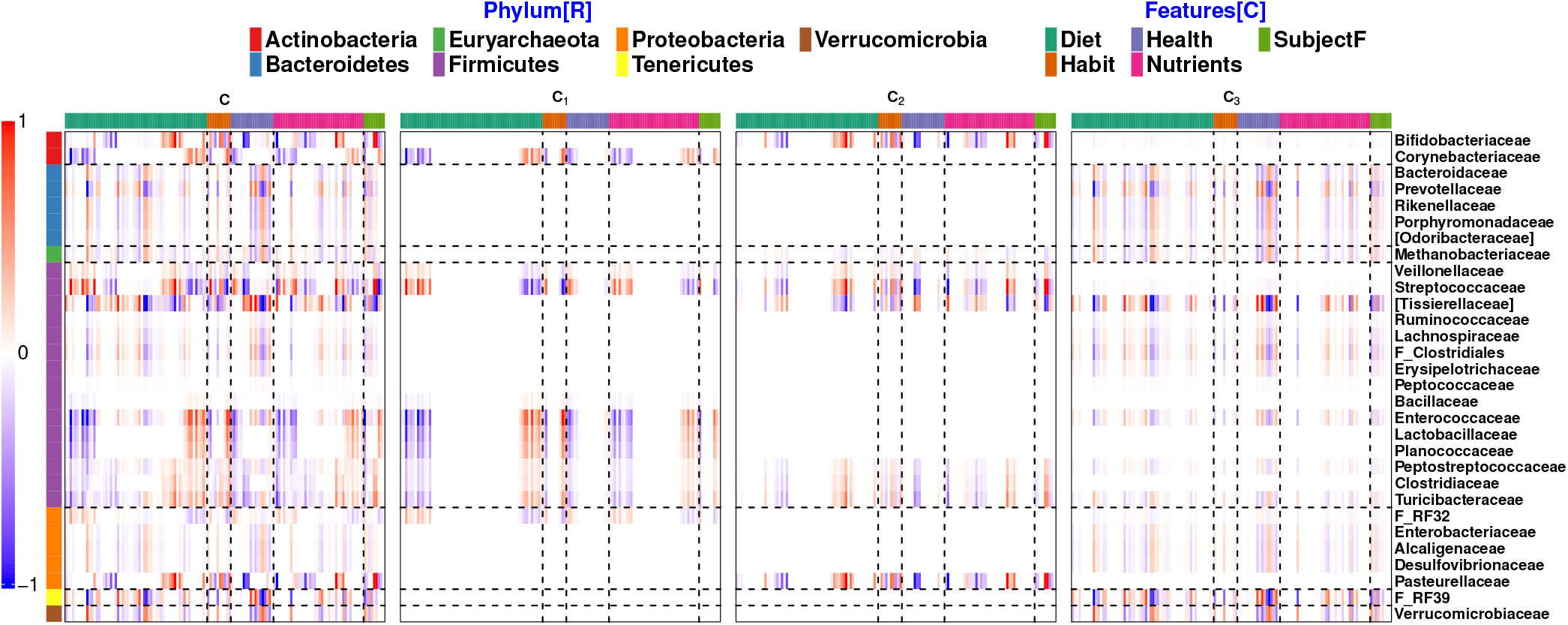
Application – AGP: The sparse estimate of the selected rows and columns of the coefficient matrix **Ĉ** with its corresponding unit-rank components using NB-FAR. Based on the phylum of the taxon, horizontal lines separate the response into 7 categories: Actinobacteria, Bacteroidetes, Euryarchaeota, Firmicutes, Proteobacteria, Tenericutes and Verrucomicrobia (top to bottom). Based on the type of the covariates, vertical lines (left to right) separate the selected predictors into five categories: diet, habit, health, nutrients and subject features.

For instance, let us consider the top-left sub-matrix of **C** which relates the two Actinobacteria *Bifidobacteriaceae* and *Corynebacteriaceae* (red) to diet covariates (green). Here, we observe an almost disjoint association pattern of positive and negative factors with diet covariates for these two families. This pattern arises from the first two latent factor **C**_1_ and **C**_2_. There, we observe a unique non-zero association pattern of diet variables with *Corynebacteriaceae* (second row in **C**_1_) and a unique non-zero association pattern of diet variables with *Bifidobacteriaceae* (first row in **C**_2_). The third latent component does not contribute any additional associations.

We next focused on the analysis of the overall most important covariates-taxa associations. The left panel of Figure 4 shows the top 25 covariates based on the row sum 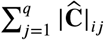 of the absolute values of the estimated coefficient matrix; the right plot shows the union of the top 10 covariates selected by each of the three unit-rank components of the estimated **C**. The color intensity in the plot reflects the effect of covariates on the abundance (red/blue for positive/negative effect). For instance, our analysis suggests that latitude (an indicator of geography) or cat ownership significantly impact the abundance of *Bifidobacteriaceae, Pasteurellaceae*, and *Streptococcaceae*. This finding is supported by several studies that also report a strong role of geography^51,52,53^ and pet ownership^54,55^ on microbial abundance patterns. Likewise, the consumption of olive oil (unsaturated fatty acids) negatively impacts the abundance of *Corynebacteriaceae* and of several families in the Firmicutes phylum while showing no influence on Bacteroidetes families. Several studies such as De Wit et al. (2012)^56^, Zhao et al. (2019)^57^ and Farras et al. (2020)^58^ have reported associations between olive oil intake and a reduction in *Firmicutes/Bacteroidetes* ratio, consistent with the observations here. The NB-FAR model also suggests that *Bifidobacteriaceae* abundances are positively associated with grain intake (*vioscreen_hei_grain:* Healthy Eating Index (HEI) Score of total grain), as previously observed^59^. Finally, we also observed several associations between an underlying medical condition and microbial taxa. For instance, attention deficit hyperactivity disorder (ADHD) (last column in left panel, *add_adhd*) negatively impacts the abundance of *Streptococcaceae, Bifidobacteriaceae*, and *Pasteurellaceae*. A similar observation about *Streptococcaceae* has been reported in a separate study on children with Autism Spectrum Disorder (showing signs of ADHD)^60^.

**FIGURE 4.**
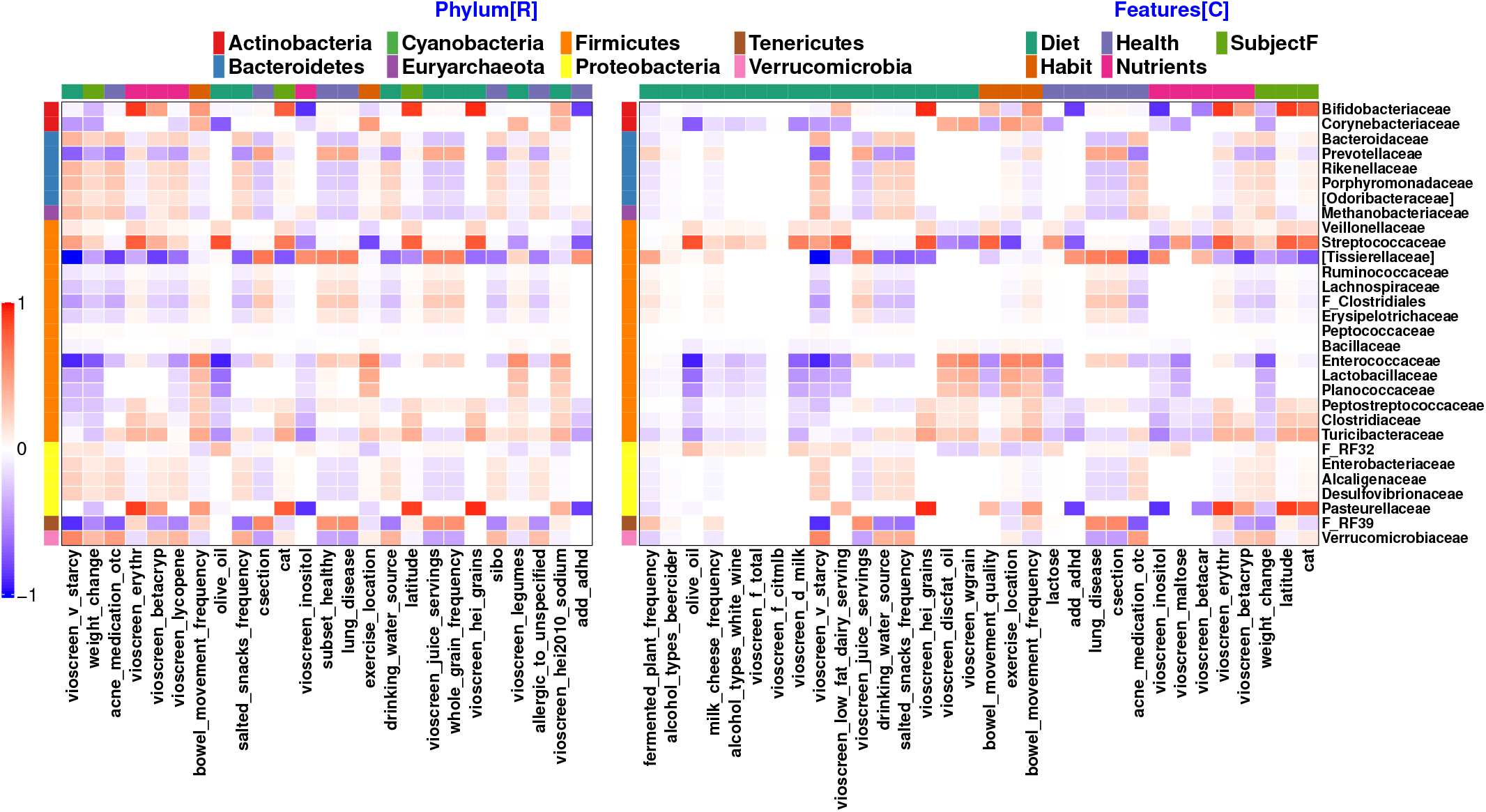
Application – AGP: Plots show the selected rows and columns of the coefficient matrix **C** based on the estimated effect size. The left plot selects the top 25 covariates based on the row sum of the absolute values of the estimated coefficient matrix,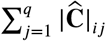.The right plot shows the union of the top 10 covariates selected by each of the three unit-rank components of the estimated **C**.

The orthogonality property of the three latent factors also allows a unique factor-by-factor analysis of the estimated associations. Since each unit-rank component estimate divides response-predictor pairs into two groups, i.e., positive and negative, we can cluster each component into exactly four quadrants, essentially enabling disentangled bi-clustering of microbial families and host-associated covariates. Figure 5 illustrates these biclusters for each of the three unit-rank components {**C**_1_, **C**_2_, **C**_3_}. For instance, in the **C**_1_ estimate plot, the upper left quadrant shows the negative associations between subsets of covariates and subsets of families. The covariate olive oil intake (column A1) is significantly negatively associated with seven out of the eight taxa, including *Corynebacteriaceae* and *Enterococcaceae*. On the other hand, the covariate exercise location (column A3) is positively associated with these (and other) taxa. Similarly, from the **C**_2_ estimate, we observed that the taxa *Bifidobacteriaceae, Pasteurellaceae* and *Streptococcaceae* are positively associated with latitude/geography, Erythritol intake, and grain (columns B1-B3), and negatively associated with inositol level (nutrients) and ADHD (columns B4-B5). Finally, from the **C**_3_ estimate, we observe that the covariates sets {starchy vegetable, acne medication, salted snack frequency, drinking water source} (columns C1,C2,C3,C6) and {cesarean birth, lung diseases, juice serving} (columns C4,C5,C7) have significantly opposite effects on the **C**_3_-associated microbial families.

**FIGURE 5.**
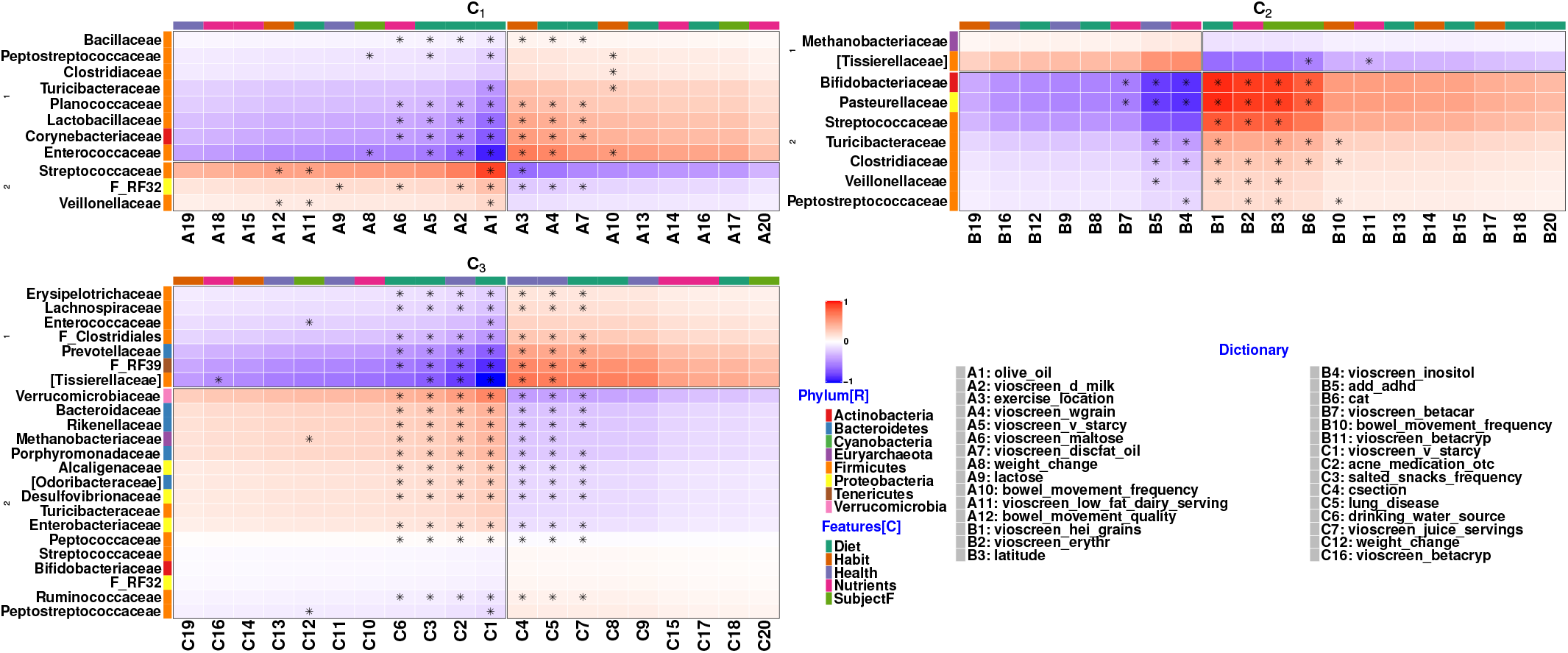
Application – AGP: Plots shows the selected rows and columns of the estimated unit-rank components, i.e., {**C**_1_, **C**_2_, **C**_3_}, of the coefficient matrix **C**. Covariates selected in the right plot of the Figure (4) are marked ⋆ in each of the three components. Each quadrant in the subplots shows the underlying associations.

Overall, these results highlight a strong influence of the host-associated features on microbial abundance pattern which should be accounted for whenever such features are available in a microbiome study.

## 6 DISCUSSION

In this contribution, we have presented two novel Negative Binomial factor regression models, NB-FAR and NB-RRR, for the analysis of microbial abundance data. The models have been tailored toward modeling overdispersed count outcome data in the context of amplicon-derived microbiome data. However, we posit that the models may prove useful in other application areas, including clinical trails ^61^, sports ^62^, and single-cell genomics^63^. The key novelty of the models is to express underlying dependencies between responses (e.g., microbial counts) and predictors (e.g., host or environmental covariates) by assuming either a dense low-rank or a co-sparse low-rank representation of the coefficient matrix. These structural assumptions appear realistic in the context of microbiome data where certain bacterial taxa are likely specialized in metabolizing specific food ingredients and hence show a concerted diet dependence. Compared to marginal approaches where each of the families is separately modeled using a Poisson or Negative Binomial regression, we have shown that our models are both computationally more efficient and simultaneously achieve better estimation performance on simulated data and the large-scale American Gut Project (AGP) data compendium. These results, in turn, challenge recent efforts in establishing host-microbiome relationships using marginal approaches^32,64^. In particular, the study by Manor et al. (2020)^64^ attempted to estimate large-scale association patterns between microbial genera and host features across thousands of participants using marginal logistic or Poisson regression. There, the authors used microbial abundances as predictors and reported the most significant association patterns with individual host covariates as outcome. In particular, they identified the genera Ys2, Ml615j–28, Coriobacteriaceae, Christensenellaceae, Mogibacteriaceae, and Oxalobacter to be significantly associated with host covariates all of which where among the few families that were *not* associated with host covariates in our analysis. These discrepancies highlight the fact that, despite reasonably large sample sizes, there still remain considerable inconsistencies across microbiome studies that require further statistical (meta-)analysis. As we have shown in our application on the AGP data, the ability of the NB-FAR model to deliver crisp bi-clustering of the underlying host-microbiome associations makes them an ideal tool to perform such future analysis and generate testable biological hypotheses.

On the statistical side, potentially fruitful extensions of the Negative Binomial factor models include the handling of excess zeros in the outcome data using, e.g., zero-inflated components or hurdle models. Since there are two types of zeros in the microbial abundance data, true (structural) zeros and experimental (measurement) zeros, one potential path forward is to identify structural zeros *a priori*^65^ and treat the remaining zeros as missing values the latter of which can already be efficiently handled by NB-FAR and NB-RRR. Furthermore, our current approach is restricted to using the *log* link function associating the mean to the linear predictor. Introducing alternative link functions that satisfy the positive mean constraint would add to the flexibility and generality of the current modeling framework. Finally, in our current framework we select the tuning parameter *λ* via K-fold cross-validation, making the parameter estimation procedure computationally intensive. This can potentially be alleviated by developing a stage-wise algorithm^66^ for parameter estimation since such a strategy has been proven to be computationally efficient in the multivariate linear regression setting with normally distributed response matrix **Y**.

Going forward, we also posit that NB-FAR and NB-RRR may serve as useful sub-routines in more complex statistical analysis workflows, including causal inference. For example, consider a typical randomized clinical trial experiment that aims at understanding the causal effect of a treatment on a phenotype of interest. In a diet intervention study, for instance, it is not unlikely that the intended direct effect on host health is mediated or confounded by the presence of certain microbes in the microbiome. While the instrumental variable (IV) approach^67^ provides a powerful framework to uncover causal effects, it requires that the instruments are strong and not confounded. A standard IV approach for continuous data estimates the parameters using two-stage least square. For the high-dimensional data problem, Lin et al. (2015)^68^ proposed a regularized two-stage framework that solves a penalized multivariate linear regression in the first stage and a Lasso problem^37^ in the second stage. We posit that this framework can be extended for the overdispersed count data by using the NB-FAR/NB-RRR methodology in the first stage when overdispersed count data serve as the independent variables.

Taken together, we believe that the introduced Negative Binomial factor regression models and their efficient implementation in the R package `nbfar` provide a useful statistical framework for analyzing overdispersed count data in medicine, biology, and other scientific disciplines.

## Supporting information

Supplementary analysis

## Abbreviations

NB-FAR: Negative Binomial co-sparse factor regression
NB-RRR: Negative Binomial reduced rank regression

## ACKNOWLEDGEMENTS

The authors would like to thank Dr. Andreas Buja for his comments and suggestions in developing the proposed procedure in the manuscript.

## DATA AVAILABILITY STATEMENT

The R code for the simulation study and an application for analyzing the microbiome data from American Gut Project is available as `simulationExamples.R`. All the supporting files are available on the GitHub page of the project https://github.com/amishra-stats/nbfar.

## Notes

https://github.com/amishra-stats/nbfar

### Competing Interest Statement

The authors have declared no competing interest.

