## Supplementary analysis for "Negative Binomial factor regression with application to microbiome data analysis"

November 28, 2021

### 1 Supplementary Materials

#### 1.1 Estimation Procedure for NB-RRR

We follow the framework of mixed-reduced-rank-regression (M-RRR) [Luo et al., 2018] to compute the parameters estimate of NB-RRR by solving the optimization problem (8). Our iterative procedure alternate between $\mathbf{C}$-step, $\boldsymbol{\beta}$-step and $\boldsymbol{\Phi}$-step to update the parameters $\mathbf{C}$, $\boldsymbol{\beta}$ and $\boldsymbol{\Phi}$, respectively, until convergence.

**C-step** For fixed $\{\boldsymbol{\beta}, \boldsymbol{\Phi}\}$, we denote $\boldsymbol{\Theta}$, a function of $\mathbf{C}$, as $\boldsymbol{\Theta}(\mathbf{C})$. Suppose differentiable $\mathcal{L}(\boldsymbol{\Theta}(\mathbf{C}), \boldsymbol{\Phi})$ is L-Lipschitz continuous gradient for some $L_c$ i.e., $\|\boldsymbol{\nabla}\mathcal{L}(\boldsymbol{\Theta}(\check{\mathbf{C}})) - \boldsymbol{\nabla}\mathcal{L}(\boldsymbol{\Theta}(\mathbf{C}))\| \leq L_c\|\check{\mathbf{C}} - \mathbf{C}\|$ for any conformable $\check{\mathbf{C}}$. The statement holds for any $\sup_{\mathbf{C}} \|\boldsymbol{\nabla}^2\mathcal{L}(\boldsymbol{\Theta}(\mathbf{C}))\| \leq L_c$. Using the result, we majorize $\mathcal{L}(\boldsymbol{\Theta}(\mathbf{C}), \boldsymbol{\Phi})$ by a convex surrogate at a given $\check{\mathbf{C}}$ as

$$\mathcal{L}(\boldsymbol{\Theta}(\mathbf{C}), \boldsymbol{\Phi}) \leq M(\mathbf{C}; \check{\mathbf{C}}) = \mathcal{L}(\boldsymbol{\Theta}(\check{\mathbf{C}})), \boldsymbol{\Phi}) + \mathrm{tr}(\boldsymbol{\nabla}\mathcal{L}(\boldsymbol{\Theta}(\check{\mathbf{C}}), \boldsymbol{\Phi}))^{\mathrm{T}}\{\mathbf{C} - \check{\mathbf{C}})\}) + L_c\|\check{\mathbf{C}} - \mathbf{C}\|^2/2. \quad (1)$$

Let us denote the current estimate of $\mathbf{C}$ by $\check{\mathbf{C}}$. We update the parameter $\mathbf{C}$ as

$$\overline{\mathbf{C}} \equiv \arg\min_{\check{\mathbf{C}}} \; M(\mathbf{C}; \check{\mathbf{C}}), \qquad \text{s.t.} \qquad \mathrm{rank}(\mathbf{C}) \leq r.$$

The unique optimal solution is given by $\overline{\mathbf{C}} = \mathbb{T}^{(r)}(\check{\mathbf{C}} - \boldsymbol{\nabla}\mathcal{L}(\boldsymbol{\Theta}(\check{\mathbf{C}}), \boldsymbol{\Phi})/L_c)$ where $\mathbb{T}^{(r)}(\mathbf{M})$ extracts $r$ singular value decomposition (SVD) components of matrix $\mathbf{M}$. In the parametric formulation (8), we relate $\boldsymbol{\Theta}(\mathbf{C})$ to the linear predictor $\boldsymbol{\eta}(\mathbf{C})$ as $g(b'(\boldsymbol{\Theta}(\mathbf{C}))) = \boldsymbol{\eta}(\mathbf{C})$. Interchangeably, we express the $\mathcal{L}(\boldsymbol{\Theta}(\mathbf{C}), \boldsymbol{\Phi})$ as function of $\boldsymbol{\eta}(\mathbf{C})$ as $\mathcal{L}(\boldsymbol{\eta}(\mathbf{C}), \boldsymbol{\Phi})$. To compute $L_c$, we define

$$\sup_{\mathbf{C}} \|\boldsymbol{\nabla}^2\mathcal{L}(\boldsymbol{\Theta}(\mathbf{C}))\| = \max_{1\leq j\leq q} \sup_{\mathbf{C}_{.j}} \|\boldsymbol{\nabla}^2\mathcal{L}(\boldsymbol{\Theta}(\mathbf{C}_{.j}))\|,$$

where $\boldsymbol{\nabla}^2\mathcal{L}(\boldsymbol{\Theta}(\mathbf{C}_{.j})) = \mathbf{X}^{\mathrm{T}}\mathrm{diag}[\boldsymbol{\nabla}^2{}_{.j}\mathcal{L}(\boldsymbol{\eta})]\mathbf{X}$ and $\boldsymbol{\nabla}^2\mathcal{L}(\boldsymbol{\eta}) = [\boldsymbol{\nabla}^2\mathcal{L}(\boldsymbol{\eta}_{ij})]_{n\times q}$ such that

$$\boldsymbol{\nabla}^2\mathcal{L}(\eta_{ij}) = (y_{ij}+1)\frac{\phi_j \exp \eta_{ij}}{(\phi_j + \exp \eta_{ij})^2} \leq \frac{y_{ij}+1}{4} \implies L_c = \max_{1\leq j\leq q} \|\mathbf{X}^{\mathrm{T}}\mathrm{diag}(\mathbf{Y}_{.j}+1)\mathbf{X}\|/2. \quad (2)$$

**$\boldsymbol{\beta}$-step:** For fixed $\mathbf{C}$ and $\boldsymbol{\Phi}$, we denote $\boldsymbol{\Theta}$, a function of $\boldsymbol{\beta}$, as $\boldsymbol{\Theta}(\boldsymbol{\beta})$. In terms of $\boldsymbol{\beta}$, suppose $\mathcal{L}(\boldsymbol{\Theta}(\boldsymbol{\beta}), \boldsymbol{\Phi})$ is L-Lipschitz continuous gradient for some $L_b$. Following the $\mathbf{C}$-step, we majorize the objective function $\mathcal{L}(\boldsymbol{\Theta}(\boldsymbol{\beta}), \boldsymbol{\Phi})$ at the current parameter $\check{\boldsymbol{\beta}}$ as

$$\mathcal{L}(\boldsymbol{\Theta}(\boldsymbol{\beta}), \boldsymbol{\Phi}) \leq K(\boldsymbol{\beta}; \check{\boldsymbol{\beta}}) = \mathcal{L}(\boldsymbol{\Theta}(\check{\boldsymbol{\beta}}), \boldsymbol{\Phi}) + \boldsymbol{\nabla}\mathcal{L}(\boldsymbol{\Theta}(\check{\boldsymbol{\beta}}), \boldsymbol{\Phi})^{\mathrm{T}}\{\boldsymbol{\beta} - \check{\boldsymbol{\beta}})\} + \frac{L_b}{2}\|\check{\boldsymbol{\beta}} - \boldsymbol{\beta}\|^2, \quad (3)$$

and update the parameter with the solution of the optimization problem

$$\overline{\boldsymbol{\beta}} \equiv \arg\min_{\boldsymbol{\beta}} \; K(\boldsymbol{\beta}; \check{\boldsymbol{\beta}}).$$

The unique optimal solution is given by $\overline{\boldsymbol{\beta}} = \check{\boldsymbol{\beta}} - \boldsymbol{\nabla}\mathcal{L}(\boldsymbol{\Theta}(\check{\boldsymbol{\beta}}), \boldsymbol{\Phi})/s_b$. Following $\mathbf{C}$-step, we compute $L_b = \max_{1\leq j\leq q} \|\mathbf{Z}^{\mathrm{T}}(\mathbf{Y}_{.j}+1)\mathbf{Z}\|/2$.

**$\boldsymbol{\Phi}$-step:** For fixed $\mathbf{C}$ and $\boldsymbol{\beta}$, we apply Newton-Raphson [R Core Team, 2019] to update the current estimate of $\boldsymbol{\Phi}$ such that the negative log-likelihood function $\mathcal{L}(\boldsymbol{\Theta}, \boldsymbol{\Phi})$ decreases; see Section 1.5 for more details on the update rule.

### 1.2 Proof of Theorem 1

We prove the monotone decreasing likelihood property of Algorithm 1 estimating parameters of the NB-RRR. Let us denote the parameters set in the $t$th iteration by $\mathbb{L}^{(t)} = (\mathbf{C}^{(t)}, \boldsymbol{\beta}^{(t)}, \Phi^{(t)})$.

**C-step** With $\{\boldsymbol{\beta}^{(t)}, \Phi^{(t)}\}$ fixed, we denote the likelihood as function of $\mathbf{C}$ by $\mathcal{L}(\boldsymbol{\Theta}(\mathbf{C}), \boldsymbol{\Phi}^{(t)})$. Using the result in equation (1), for $t$th iteration, we have

$$\mathcal{L}(\boldsymbol{\Theta}(\mathbf{C}), \boldsymbol{\Phi}^{(t)}) \leq M(\mathbf{C}; \mathbf{C}^{(t)}) \qquad \forall \qquad \mathbf{C} \in \mathbb{R}^{p\times q}.$$

Let us denote the optimal solution minimizing the surrogate function $M(\mathbf{C}; \mathbf{C}^{(t)})$ in the $\mathbf{C}$-step by $\mathbf{C}^{(t+1)}$. For $\mathbf{C}^{(t+1)}$, we write the above result as

$$\mathcal{L}(\boldsymbol{\Theta}(\mathbf{C}^{(t+1)}), \boldsymbol{\Phi}^{(t)}) \leq M(\mathbf{C}^{(t+1)}; \mathbf{C}^{(t)}) \leq M(\mathbf{C}^{(t)}; \mathbf{C}^{(t)}) = \mathcal{L}(\boldsymbol{\Theta}(\mathbf{C}^{(t)}), \boldsymbol{\Phi}^{(t)}).$$

**$\boldsymbol{\beta}$-step** With $\{\mathbf{C}^{(t+1)}, \Phi^{(t)}\}$ fixed, we denote the likelihood as function of $\boldsymbol{\beta}$ given by $\mathcal{L}(\boldsymbol{\Theta}(\boldsymbol{\beta}), \boldsymbol{\Phi}^{(t)})$. Using the result in equation (3), for $t$th iteration, we have

$$\mathcal{L}(\boldsymbol{\Theta}(\boldsymbol{\beta}), \boldsymbol{\Phi}^{(t)}) \leq K(\boldsymbol{\beta}; \boldsymbol{\beta}^{(t)}) \qquad \forall \qquad \boldsymbol{\beta} \in \mathbb{R}^{c\times q}.$$

Let us denote the optimal solution minimizing the surrogate function $K(\boldsymbol{\beta}; \boldsymbol{\beta}^{(t)})$ in the $\boldsymbol{\beta}$-step by $\boldsymbol{\beta}^{(t+1)}$. For $\boldsymbol{\beta}^{(t+1)}$, we write the above result as

$$\mathcal{L}(\boldsymbol{\Theta}(\boldsymbol{\beta}^{(t+1)}), \boldsymbol{\Phi}^{(t)}) \leq K(\boldsymbol{\beta}^{(t+1)}; \boldsymbol{\beta}^{(t)}) \leq K(\boldsymbol{\beta}^{(t)}; \boldsymbol{\beta}^{(t)}) = \mathcal{L}(\boldsymbol{\Theta}(\boldsymbol{\beta}^{(t)}), \boldsymbol{\Phi}^{(t)}).$$

Finally, in $\boldsymbol{\Phi}$-step, with fixed $\mathbf{C}^{(t+1)}$ and $\boldsymbol{\beta}^{(t+1)}$, we minimize the negative log-likelihood function $\mathcal{L}(\boldsymbol{\Theta}, \boldsymbol{\Phi})$ with respect to $\boldsymbol{\Phi}$ using Newton-Raphson [R Core Team, 2019]. The approach guarantees to have non-increasing likelihood function $\mathcal{L}(\boldsymbol{\Theta}, \boldsymbol{\Phi})$.

Combining the result of the non-increasing negative log-likelihood function $\mathcal{L}(\boldsymbol{\Theta}, \boldsymbol{\Phi})$ in $\mathbf{C}$-step, $\boldsymbol{\beta}$-step and $\boldsymbol{\Phi}$-step, we prove the monotone decreasing likelihood property of Algorithm 1.

### 1.3 Estimation Procedure for Solving NB-CURE Problem

We follow the framework of generalize co-sparse factor regression (GO-FAR) [Mishra et al., 2020] to compute the parameters estimate of NB-CURE by solving the optimization problem (11). Our iterative procedure alternate between $\mathbf{u}$-step, $\mathbf{v}$-step, $\boldsymbol{\beta}$-step and $\boldsymbol{\Phi}$-step to update the parameters in blocks of $(\mathbf{u}, d)$, $(\mathbf{v}, d)$, $\boldsymbol{\beta}$ and $\boldsymbol{\Phi}$, respectively, until convergence.

**u-step:** For fixed $\{\mathbf{v}, \boldsymbol{\beta}, \boldsymbol{\Phi}\}$ with $\mathbf{v}^{\mathrm{T}}\mathbf{v} = 1$, we express the $\boldsymbol{\Theta}$ as a function of the product variable $\check{\mathbf{u}} = d\mathbf{u}$, denoted by $\boldsymbol{\Theta}(\check{\mathbf{u}}\mathbf{v}^{\mathrm{T}})$, and the objective function in (11) as $F_\lambda(\check{\mathbf{u}}, \mathbf{v}, \boldsymbol{\beta}, \boldsymbol{\Phi})$. In terms of $\check{\mathbf{u}}$, let us assume that the $\mathcal{L}(\boldsymbol{\Theta}(\check{\mathbf{u}}\mathbf{v}^{\mathrm{T}}), \boldsymbol{\Phi})$ is L-Lipschitz continuous gradient for some $L_u$. Following the $\mathbf{C}$-step in Section 1.1 for NB-RRR, we majorize the objective function (11) as

$$F_\lambda(\mathbf{a}, \mathbf{v}, \boldsymbol{\beta}, \boldsymbol{\Phi}) = \mathcal{L}(\boldsymbol{\Theta}(\mathbf{a}\mathbf{v}^{\mathrm{T}}), \boldsymbol{\Phi}) + \rho_\lambda(\mathbf{a}\mathbf{v}^{\mathrm{T}}; \mathbf{W}) \leq G_\lambda(\mathbf{a}; \check{\mathbf{u}}) \qquad \forall \qquad \mathbf{a} \in \mathbb{R}^p, \tag{4}$$

$$\text{where} \quad G_\lambda(\mathbf{a}; \check{\mathbf{u}}) = \mathcal{L}(\boldsymbol{\Theta}(\check{\mathbf{u}}\mathbf{v}^{\mathrm{T}}), \boldsymbol{\Phi}) + \boldsymbol{\nabla}\mathcal{L}(\boldsymbol{\Theta}(\check{\mathbf{u}}\mathbf{v}^{\mathrm{T}}), \boldsymbol{\Phi}))^{\mathrm{T}}(\mathbf{a} - \check{\mathbf{u}}) + \frac{L_u}{2}\|\check{\mathbf{u}} - \mathbf{a}\|^2 + \rho_{\lambda)}(\mathbf{a}\mathbf{v}^{\mathrm{T}}; \mathbf{W}),$$

and solve the optimization problem

$$\widehat{\mathbf{u}} \equiv \arg\min_{\mathbf{a}} \; G_\lambda(\mathbf{a}; \check{\mathbf{u}}),$$

to update $\check{\mathbf{u}}$. Following Zou and Hastie [2005], the unique optimal solution is given by

$$\widehat{\mathbf{u}} = \frac{\mathbf{S}(\check{\mathbf{u}} - \boldsymbol{\nabla}\mathcal{L}(\boldsymbol{\Theta}(\check{\mathbf{u}}\mathbf{v}^{\mathrm{T}}), \boldsymbol{\Phi})/L_u; \alpha\lambda\mathbf{v}^{\mathrm{T}}\mathbf{w}^{(v)}w^{(d)}\mathbf{w}^{(u)}/L_u)}{1 + 2\lambda(1-\alpha)\|\mathbf{v}\|_2^2/L_u}, \tag{5}$$

where $\mathbf{S}(\mathbf{t}; \tilde{\lambda}) = \mathrm{sign}(\mathbf{t})(|\mathbf{t}| - \tilde{\lambda})_+$ is the elementwise soft-thresholding operator on any $\mathbf{t} \in \mathbb{R}^p$. We utilize the equality constraint in equation (11) to recover the estimate of $(d, \mathbf{u})$ from $\widehat{\mathbf{u}}$. To compute $L_u$, we derive $\frac{\partial \mathcal{L}}{\partial \check{\mathbf{u}}} = \mathbf{X}^{\mathrm{T}}\boldsymbol{\nabla}\mathcal{L}(\boldsymbol{\eta})\mathbf{v} = \sum_{i=1}^n \mathbf{x}_i \sum_{k=1}^q \boldsymbol{\nabla}\mathcal{L}(\eta_{ik})v_k$ and $\frac{\partial^2 \mathcal{L}}{\partial \check{\mathbf{u}}\partial \check{\mathbf{u}}^{\mathrm{T}}} = \sum_{i=1}^n \mathbf{x}_i \Big(\sum_{k=1}^q \boldsymbol{\nabla}^2\mathcal{L}(\eta_{ik})v_k^2\Big)\mathbf{x}_i^{\mathrm{T}}$ and utilize the result in (2) to upper bound the $\sup_{\check{\mathbf{u}}} \|\frac{\partial^2 \mathcal{L}}{\partial \check{\mathbf{u}}\partial \check{\mathbf{u}}^{\mathrm{T}}}\|$ as

$$L_u = \|\sum_{i=1}^n \mathbf{x}_i \Big(\sum_{k=1}^q (y_{ik}+1)v_k^2\Big)\mathbf{x}_i^{\mathrm{T}}\|/2 = \|\mathbf{X}^{\mathrm{T}}\mathbf{X} + \sum_{i=1}^n \mathbf{x}_i \Big(\sum_{k=1}^q y_{ik}v_k^2\Big)\mathbf{x}_i^{\mathrm{T}}\|/2.$$

**v-step** For fixed $\{\mathbf{u}, \boldsymbol{\beta}, \boldsymbol{\Phi}\}$, we express $\boldsymbol{\Theta}$ as a function of the product variable $\check{\mathbf{v}} = d\mathbf{v}$, denoted by $\boldsymbol{\Theta}(\mathbf{u}\check{\mathbf{v}}^{\mathrm{T}})$, and the objective function in (11) as $F_\lambda(\mathbf{u}, \check{\mathbf{v}}, \boldsymbol{\beta}, \boldsymbol{\Phi})$. In terms of $\check{\mathbf{v}}$, suppose $\mathcal{L}(\boldsymbol{\Theta}(\mathbf{u}\check{\mathbf{v}}^{\mathrm{T}}), \boldsymbol{\Phi})$ is L-Lipschitz continuous gradient for some $L_v$. Again we majorize the objective function (11) as

$$F_\lambda(\mathbf{u}, \mathbf{b}, \boldsymbol{\beta}, \boldsymbol{\Phi}) = \mathcal{L}(\boldsymbol{\Theta}(\mathbf{u}\mathbf{b}^{\mathrm{T}}), \boldsymbol{\Phi}) + \rho_\lambda(\mathbf{u}\mathbf{b}^{\mathrm{T}}; \mathbf{W}) \leq H_\lambda(\mathbf{b}; \check{\mathbf{v}}) \qquad \forall \qquad \mathbf{b} \in \mathbb{R}^q, \tag{6}$$

$$\text{where} \quad H_\lambda(\mathbf{b}; \check{\mathbf{v}}) = \mathcal{L}(\boldsymbol{\Theta}(\mathbf{u}\check{\mathbf{v}}^{\mathrm{T}}), \boldsymbol{\Phi}) + \boldsymbol{\nabla}\mathcal{L}(\boldsymbol{\Theta}(\mathbf{u}\check{\mathbf{v}}^{\mathrm{T}}), \boldsymbol{\Phi})^{\mathrm{T}}(\mathbf{b} - \check{\mathbf{v}}) + \frac{L_v}{2}\|\check{\mathbf{v}} - \mathbf{b}\|^2 + \rho_{\lambda)}(\mathbf{u}\mathbf{b}^{\mathrm{T}}; \mathbf{W}),$$

and solve the optimization problem

$$\widehat{\mathbf{v}} \equiv \arg\min_{\mathbf{b}} H_\lambda(\mathbf{b}; \check{\mathbf{v}}),$$

to update $\check{\mathbf{v}}$. Following the $\mathbf{u}$-step, the unique optimal solution is given by

$$\widehat{\mathbf{v}} = \frac{\mathbf{S}(\check{\mathbf{v}} - \boldsymbol{\nabla}\mathcal{L}(\boldsymbol{\Theta}(\mathbf{u}\check{\mathbf{v}}^{\mathrm{T}}), \boldsymbol{\Phi})/L_v; \alpha\lambda\mathbf{u}^{\mathrm{T}}\mathbf{w}^{(u)}w^{(d)}\mathbf{w}^{(v)}/L_v}{1 + 2\lambda(1-\alpha)\|\mathbf{u}\|_2^2/L_v}. \tag{7}$$

Again, we utilize the equality constraint (11) to recover the estimate of $(d, \mathbf{v})$ from $\widehat{\mathbf{v}}$. To compute $L_v$, we derive $\frac{\partial \mathcal{L}}{\partial \mathbf{v}} = \Big(\frac{\partial \mathcal{L}}{\partial \boldsymbol{\eta}}\Big)^{\mathrm{T}}\mathbf{X}\mathbf{u} = \sum_{i=1}^n \mathbf{x}_i^{\mathrm{T}}\mathbf{u}\boldsymbol{\nabla}\mathcal{L}(\boldsymbol{\eta}_{i.})$ and $\frac{\partial^2 \mathcal{L}}{\partial \mathbf{v}\partial \mathbf{v}^{\mathrm{T}}} = \mathrm{diag}[\mathbf{u}^{\mathrm{T}}\mathbf{X}^{\mathrm{T}}\mathrm{diag}(\boldsymbol{\nabla}^2\mathcal{L}(\boldsymbol{\eta}_{.1}))\mathbf{X}\mathbf{u}, \ldots, \mathbf{u}^{\mathrm{T}}\mathbf{X}^{\mathrm{T}}\mathrm{diag}(\boldsymbol{\nabla}^2\mathcal{L}(\boldsymbol{\eta}_{.q}))\mathbf{X}\mathbf{u}]$. We utilize the result in (2) and upper bound $\sup_{\check{\mathbf{v}}} \|\frac{\partial^2 \mathcal{L}}{\partial \check{\mathbf{v}}\partial \check{\mathbf{v}}^{\mathrm{T}}}\|$ as

$$L_v = \max_{1\leq j\leq q} \mathbf{u}^{\mathrm{T}}\mathbf{X}^{\mathrm{T}}\mathrm{diag}(\boldsymbol{\nabla}^2\mathcal{L}(\boldsymbol{\eta}_{.j}))\mathbf{X}\mathbf{u}/2 = \max_{1\leq j\leq q} \mathbf{u}^{\mathrm{T}}\mathbf{X}^{\mathrm{T}}\mathrm{diag}[\mathbf{Y}_{.j} + 1]\mathbf{X}\mathbf{u}/2.$$

**$\boldsymbol{\beta}$-step:** For fixed $\mathbf{C} = d\mathbf{u}\mathbf{v}^{\mathrm{T}}$ and $\boldsymbol{\Phi}$, we follow $\boldsymbol{\beta}$-step update presented in Section 1.1 of NB-RRR to update the parameter $\boldsymbol{\beta}$.

**$\boldsymbol{\Phi}$-step:** For fixed $\mathbf{C}$ and $\boldsymbol{\beta}$, we apply Newton-Raphson [R Core Team, 2019] to update the current estimate of $\boldsymbol{\Phi}$ such that the negative log-likelihood function $\mathcal{L}(\boldsymbol{\Theta}, \boldsymbol{\Phi})$ decreases; see Section 1.5 for more details on the update rule.

### 1.4 Proof of Theorem 2

We prove the monotone decreasing likelihood property of Algorithm 3 solving the NB-CURE optimization problem. Let us denote the parameters set in the $t$th iteration by $\mathrm{L}^{(t)} = (d^{(t)}, \mathbf{u}^{(t)}, \mathbf{v}^{(t)}, \boldsymbol{\beta}^{(t)}, \Phi^{(t)})$.

**u-step** With $\{\mathbf{v}^{(t)}, \boldsymbol{\beta}^{(t)}, \Phi^{(t)}\}$ fixed, we update the block variables $\{d^{(t)}, \mathbf{u}^{(t)}\}$ as $\{\tilde{d}^{(t+1)}, \mathbf{u}^{(t+1)}\}$ using equation (5). Using the result in (4), we have

$$F_\lambda(\tilde{d}^{(t+1)}, \mathbf{u}^{(t+1)}, \mathbf{v}^{(t)}, \boldsymbol{\beta}^{(t)}, \Phi^{(t)}) \leq G_\lambda(\check{\mathbf{u}}^{(t+1)}; \check{\mathbf{u}}^{(t)}) \leq G_\lambda(\check{\mathbf{u}}^{(t)}; \check{\mathbf{u}}^{(t)}) = F_\lambda(d^{(t)}, \mathbf{u}^{(t)}, \mathbf{v}^{(t)}, \boldsymbol{\beta}^{(t)}, \Phi^{(t)}),$$

where $\check{\mathbf{u}}^{(t)} = d^{(t)}\mathbf{u}^{(t)}$ and $\check{\mathbf{u}}^{(t+1)} = \tilde{d}^{(t+1)}\mathbf{u}^{(t+1)}$.

**v-step** With $\{\mathbf{u}^{(t+1)}, \boldsymbol{\beta}^{(t)}, \Phi^{(t)}\}$ fixed, we update the block variables $\{\tilde{d}^{(t+1)}, \mathbf{v}^{(t)}\}$ as $\{d^{(t+1)}, \mathbf{u}^{(t+1)}\}$ using equation (7). Using the result in (6), we have

$$F_\lambda(d^{(t+1)}, \mathbf{u}^{(t+1)}, \mathbf{v}^{(t+1)}, \boldsymbol{\beta}^{(t)}, \Phi^{(t)}) \leq H_\lambda(\check{\mathbf{v}}^{(t+1)}; \check{\mathbf{v}}^{(t)}) \leq H_\lambda(\check{\mathbf{v}}^{(t)}; \check{\mathbf{v}}^{(t)}) = F_\lambda(\tilde{d}^{(t+1)}, \mathbf{u}^{(t+1)}, \mathbf{v}^{(t)}, \boldsymbol{\beta}^{(t)}, \Phi^{(t)}),$$

where $\check{\mathbf{v}}^{(t)} = \tilde{d}^{(t+1)}\mathbf{v}^{(t)}$ and $\check{\mathbf{v}}^{(t+1)} = d^{(t+1)}\mathbf{v}^{(t+1)}$.

**$\boldsymbol{\beta}$-step** With $\{\mathbf{u}^{(t+1)}, d^{(t+1)}, \mathbf{v}^{(t+1)}, \Phi^{(t)}\}$ fixed, we update $\boldsymbol{\beta}^{(t)}$ as $\boldsymbol{\beta}^{(t+1)}$ by following $\boldsymbol{\beta}$-step update presented in Section 1.1 of NB-RRR. On extending the non-decreasing result of $\boldsymbol{\beta}$-step update presented in Section 1.2, we have

$$F_\lambda(d^{(t+1)}, \mathbf{u}^{(t+1)}, \mathbf{v}^{(t+1)}, \boldsymbol{\beta}^{(t+1)}, \Phi^{(t)}) \leq F_\lambda(d^{(t+1)}, \mathbf{u}^{(t+1)}, \mathbf{v}^{(t+1)}, \boldsymbol{\beta}^{(t)}, \Phi^{(t)}).$$

Finally, in $\boldsymbol{\Phi}$-step, with fixed $\mathbf{C}^{(t+1)} = d^{(t+1)}\mathbf{u}^{(t+1)}\mathbf{v}^{(t+1)\mathrm{T}}$ and $\boldsymbol{\beta}^{(t+1)}$, we minimize the negative log-likelihood function $\mathcal{L}(\boldsymbol{\Theta}, \boldsymbol{\Phi})$ with respect to $\boldsymbol{\Phi}$ using Newton-Raphson [R Core Team, 2019]. The approach guarantees to have non-increasing likelihood function $\mathcal{L}(\boldsymbol{\Theta}, \boldsymbol{\Phi})$.

Combining the result of the non-increasing negative log-likelihood function $\mathcal{L}(\boldsymbol{\Theta}, \boldsymbol{\Phi})$ in **u**-step, **v**-step, $\boldsymbol{\beta}$-step and $\boldsymbol{\Phi}$-step, we prove the monotone decreasing likelihood property of Algorithm 3.

### 1.5 Newton-Raphson to update $\boldsymbol{\Phi}$:

For fixed $\mathbf{C}$ and $\boldsymbol{\beta}$, we have the mean parameter matrix as $\boldsymbol{\mu} = g^{-1}(\mathbf{O} + \mathbf{X}\mathbf{C} + \mathbf{Z}\boldsymbol{\beta})$. Following [Ismail and Jemain, 2007], we reparametrize $\mathcal{L}(\boldsymbol{\Theta}, \boldsymbol{\Phi})$ in terms of the inverse of the shape parameter by replacing $\phi_j = 1/\alpha_j$ for $j = 1, \ldots, q$ and write the likelihood function as

$$\mathcal{L}(\boldsymbol{\Theta}, \boldsymbol{\alpha}) = \sum_{i=1}^{n} \left[ \Big( \sum_{k=0}^{y_{ij}-1} \log \frac{k\alpha_j + 1}{\alpha_j} \Big) + y_{ij} \log \alpha_j - \Big( y_{ij} + \frac{1}{\alpha_j} \Big) \log(1 + \mu_{ij}\alpha_j) \right],$$

Table S1: Simulation: model evaluation based on 100 replications using various performance measures (standard deviations are shown in parentheses) in case of p = 100 with negative binomial responses.

| | M% | Er($\mathbf{C}$) | Er($\mathbf{\Theta}$) | FPR | FNR | R% | r | Er($\Phi$) | Time(s) |
|---|---|---|---|---|---|---|---|---|---|
| NB-FAR | 0 | 3.89 (0.97) | 16.56 (2.07) | 9.74 (4.00) | 0.00 (0.00) | 0.00 (0.00) | 3.00 (0.00) | 0.13 (0.08) | 157.66 (22.71) |
| NB-RRR | 0 | 19.67 (2.18) | 31.07 (1.39) | 100.00 (0.00) | 0.00 (0.00) | 0.00 (0.00) | 3.00 (0.00) | 0.36 (0.10) | 48.49 (6.61) |
| GO-FAR | 0 | 27.00 (7.01) | 43.62 (5.60) | 6.88 (5.32) | 51.47 (29.14) | 0.61 (1.83) | 3.30 (0.68) | 1.37 (0.00) | 31.95 (4.74) |
| NB-GLM | 0 | 237.57 (12.47) | 93.87 (2.34) | 100.00 (0.00) | 0.00 (0.00) | 51.72 (2.10) | 29.29 (0.72) | 4.54 (0.41) | 74.62 (21.17) |
| NB-FAR | 20 | 5.19 (1.73) | 18.53 (2.49) | 10.84 (4.66) | 0.59 (1.03) | 0.00 (0.00) | 3.00 (0.00) | 0.15 (0.08) | 162.06 (21.45) |
| NB-RRR | 20 | 22.22 (2.99) | 33.04 (1.48) | 100.00 (0.00) | 0.00 (0.00) | 0.00 (0.00) | 3.00 (0.00) | 0.43 (0.10) | 49.63 (7.10) |
| GO-FAR | 20 | 29.66 (8.04) | 46.98 (7.82) | 6.86 (5.67) | 50.26 (30.17) | 0.00 (0.00) | 3.00 (0.00) | 1.37 (0.00) | 35.01 (5.64) |
| NB-GLM | 20 | 294.11 (16.61) | 103.86 (2.31) | 99.87 (0.32) | 0.00 (0.00) | 55.90 (2.05) | 29.06 (1.09) | 6.04 (0.62) | 125.09 (59.07) |

where $\boldsymbol{\alpha} = \operatorname{diag}(\alpha_1, \ldots, \alpha_q)$. Based on the Newton-Raphson method, we update the current estimate $\alpha_j$ as $\hat{\alpha}_j = \alpha_j - \left(\frac{\partial^2 \mathcal{L}}{\partial \alpha_j^2}\right)^{-1} \frac{\partial \mathcal{L}}{\partial \alpha_j}$. We write the first derivative and second derivative of $\mathcal{L}(\boldsymbol{\Theta}, \boldsymbol{\alpha})$ with respect to any $\alpha_j$ as

$$
\begin{aligned}
\frac{\partial \mathcal{L}(\boldsymbol{\Theta}, \boldsymbol{\alpha})}{\partial \alpha_j} &= \sum_{i=1}^{n} \left[ \Big( \sum_{k=0}^{y_{ij}-1} \frac{k}{k\alpha_j + 1} \Big) - \frac{\mu_{ij}}{\alpha_j} \frac{y_{ij}\alpha_j + 1}{\mu_{ij}\alpha_j + 1} + \frac{1}{\alpha_j^2} \log\Big(1 + \mu_{ij}\alpha_j\Big) \right] \\
\frac{\partial^2 \mathcal{L}(\boldsymbol{\Theta}, \boldsymbol{\alpha})}{\partial \alpha_j^2} &= \sum_{i=1}^{n} \left[ \Big( \sum_{k=0}^{y_{ij}-1} \frac{-k^2}{(k\alpha_j + 1)^2} \Big) - \frac{2}{\alpha_j^3} \log\Big(1 + \mu_{ij}\alpha_j\Big) + \frac{\mu_{ij}[2 + \mu_{ij}\alpha_j(3 + y_{ij}\alpha_j)]}{\alpha_j^2(\mu_{ij}\alpha_j + 1)^2} \right].
\end{aligned}
$$

#### 1.6 Some partial derivative of $\mathcal{L}(\boldsymbol{\Theta}, \boldsymbol{\Phi})$ for fixed $\boldsymbol{\Phi}$

For the ease of notation, we write $\mathcal{L}(\boldsymbol{\Theta}, \boldsymbol{\Phi})$ as $\mathcal{L}(\boldsymbol{\Theta})$. Here $\boldsymbol{\Theta}$ is linked to linear predictor $\boldsymbol{\eta}$ as $g(b'(\boldsymbol{\Theta})) = \boldsymbol{\eta}$ and

$$
\begin{aligned}
&\mathcal{L}(\boldsymbol{\Theta}) = -\operatorname{tr}(\mathbf{Y}^{\mathrm{T}}\boldsymbol{\Theta}) + \operatorname{tr}(\mathbf{J}^{\mathrm{T}}\mathbf{B}(\boldsymbol{\Theta})) + \text{const}, \\
&\frac{\partial \mathcal{L}(\boldsymbol{\Theta})}{\partial \theta_{ij}} = -y_{ij} + b'(\theta_{ij}) \implies \frac{\partial \mathcal{L}(\boldsymbol{\Theta})}{\partial \eta_{ij}} = (-y_{ij} + b'(\theta_{ij}))\frac{\partial \eta_{ij}}{\partial \theta_{ij}}.
\end{aligned}
$$

For $g(x) = \log x$, we have $\frac{\partial \eta_{ij}}{\partial \theta_{ij}} = \frac{\phi_j}{\phi_j + \exp \eta_{ij}}$ and implies $\frac{\partial \mathcal{L}(\boldsymbol{\Theta})}{\partial \eta_{ij}} = \phi_j \Big(\frac{-y_{ij} + \exp \eta_{ij}}{\phi_j + \exp \eta_{ij}}\Big)$. Jointly we represent

$$
\frac{\partial \mathcal{L}(\boldsymbol{\Theta})}{\partial \boldsymbol{\eta}} = -\mathbf{Y} \circ \mathbf{B}^1(\boldsymbol{\eta}) + \mathbf{B}^2(\boldsymbol{\eta}),
$$

where $\mathbf{B}^1(\boldsymbol{\eta}) = [b_1(\eta_{ij})]_{n \times q}$ with $b_1(\eta_{ij}) = \frac{\phi_j}{\phi_j + \exp \eta_{ij}}$ and $\mathbf{B}^2(\boldsymbol{\eta}) = [b_2(\eta_{ij})]_{n \times q}$ with $b_2(\eta_{ij}) = \frac{\phi_j \exp \eta_{ij}}{\phi_j + \exp \eta_{ij}}$. Then, we compute $\frac{\partial \mathcal{L}(\boldsymbol{\Theta})}{\partial \boldsymbol{\beta}} = \mathbf{Z}^{\mathrm{T}} \frac{\partial \mathcal{L}(\boldsymbol{\Theta})}{\partial \boldsymbol{\eta}}$ and $\frac{\partial \mathcal{L}(\boldsymbol{\Theta})}{\partial \mathbf{C}} = \mathbf{X}^{\mathrm{T}} \frac{\partial \mathcal{L}(\boldsymbol{\Theta})}{\partial \boldsymbol{\eta}}$. When $\mathbf{C} = d\mathbf{u}\mathbf{v}^{\mathrm{T}}$, we have $\frac{\partial \mathcal{L}(\boldsymbol{\Theta})}{\partial \mathbf{u}} = \mathbf{X}^{\mathrm{T}} \frac{\partial \mathcal{L}(\boldsymbol{\Theta})}{\partial \boldsymbol{\eta}} \mathbf{v}$ and $\frac{\partial \mathcal{L}(\boldsymbol{\Theta})}{\partial \mathbf{v}} = \Big(\frac{\partial \mathcal{L}(\boldsymbol{\Theta})}{\partial \boldsymbol{\eta}}\Big)^{\mathrm{T}} \mathbf{X}\mathbf{u}$.

#### 1.7 Simulation

We report the model performance in case of $p = 100$ in Table S1.

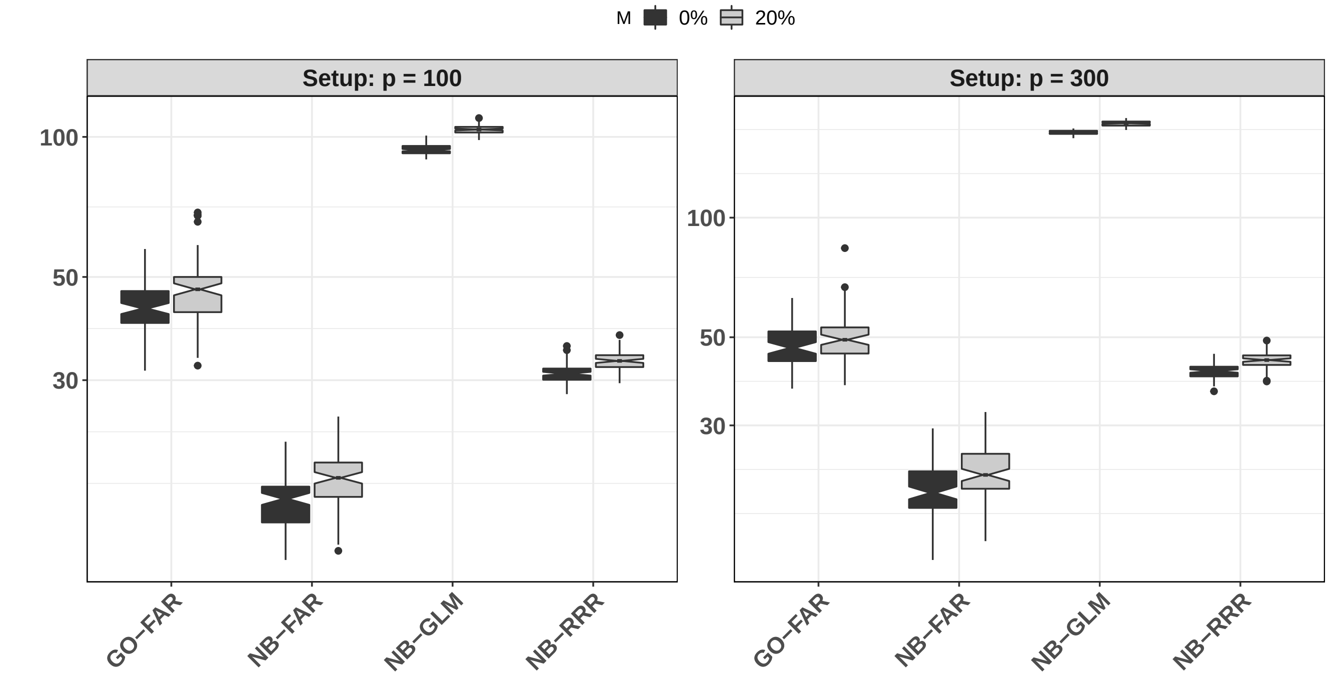


Figure S1: Simulation: notched boxplots of the estimation error $\mathrm{Er}(\boldsymbol{\eta})$ based on 100 replications in cases of p = 100 and p = 300.

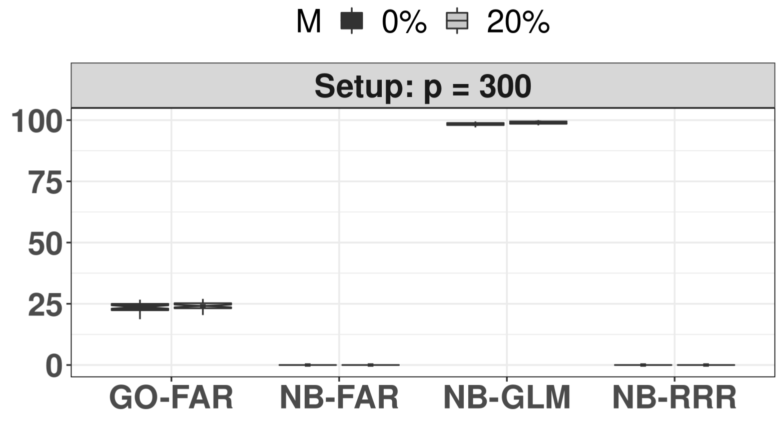


Figure S2: Simulation: notched boxplots of false positive rate, i.e., type I error, (y-axis) in a setting where true coefficient $\mathbf{C}^* = \mathbf{0}$ (null model). The analysis confirms NB-FAR's superior performance.

### 1.8 Application

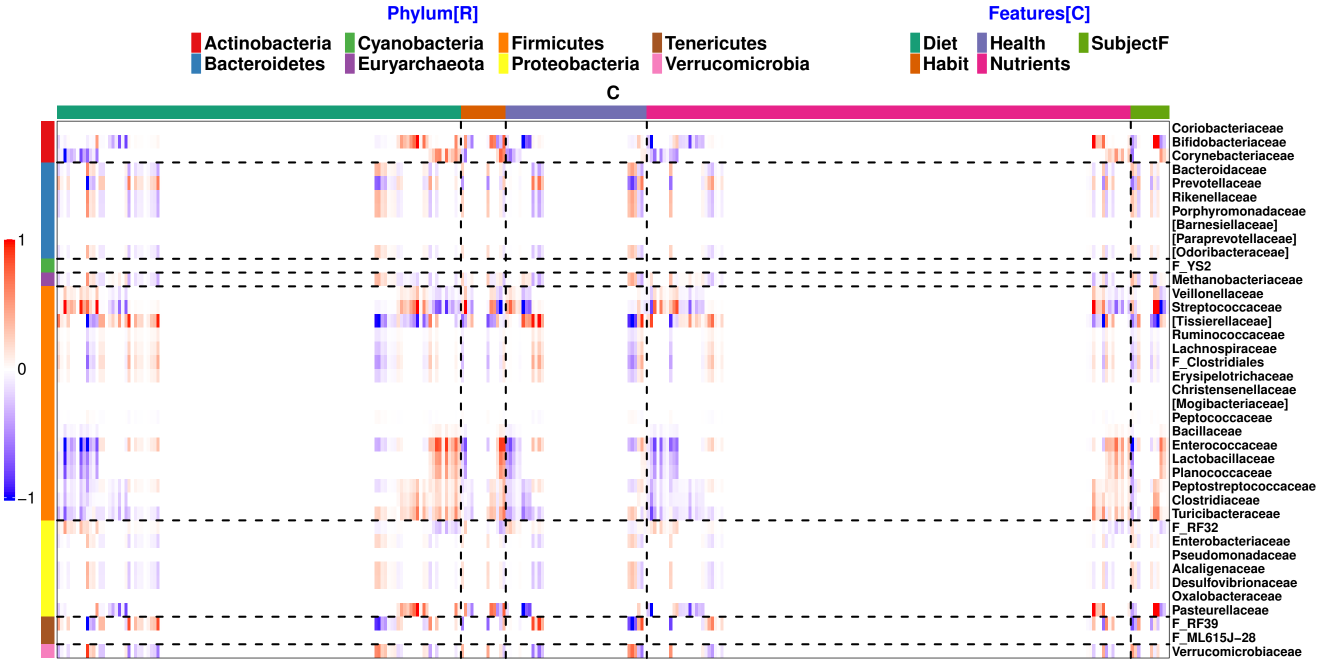


Figure S3: Application – AGP: The sparse estimate of the coefficient matrix $\widehat{\mathbf{C}}$ using NB-FAR. Based on the Phylum of the taxon, horizontal lines separate the response into 7 categories given by Actinobacteria, Bacteroidetes, Euryarchaeota, Firmicutes, Proteobacteria, Tenericutes and Verrucomicrobia (top to bottom). Based on the type of the covariates, vertical lines (left to right) separate the selected predictors into five categories: namely, diet, habit, health, nutrients and subject features.

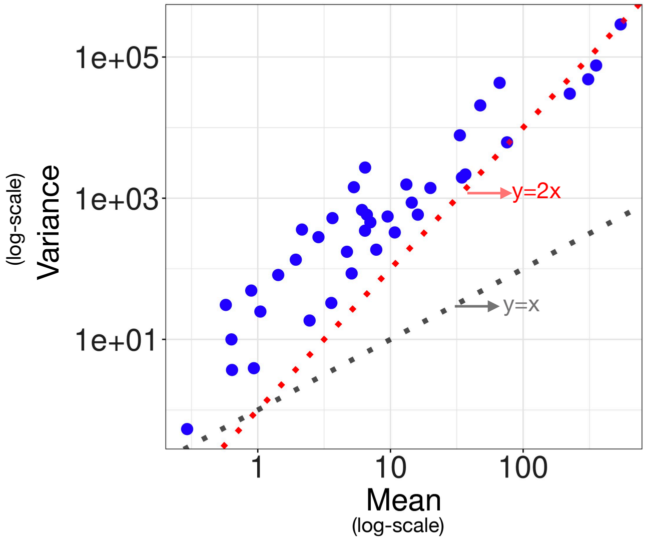


Figure S4: Application – AGP: Comparing mean and variance of the microbial abundance of each of 39 OTUs at the family level in the taxonomy to showcase overdispersed nature of the data.

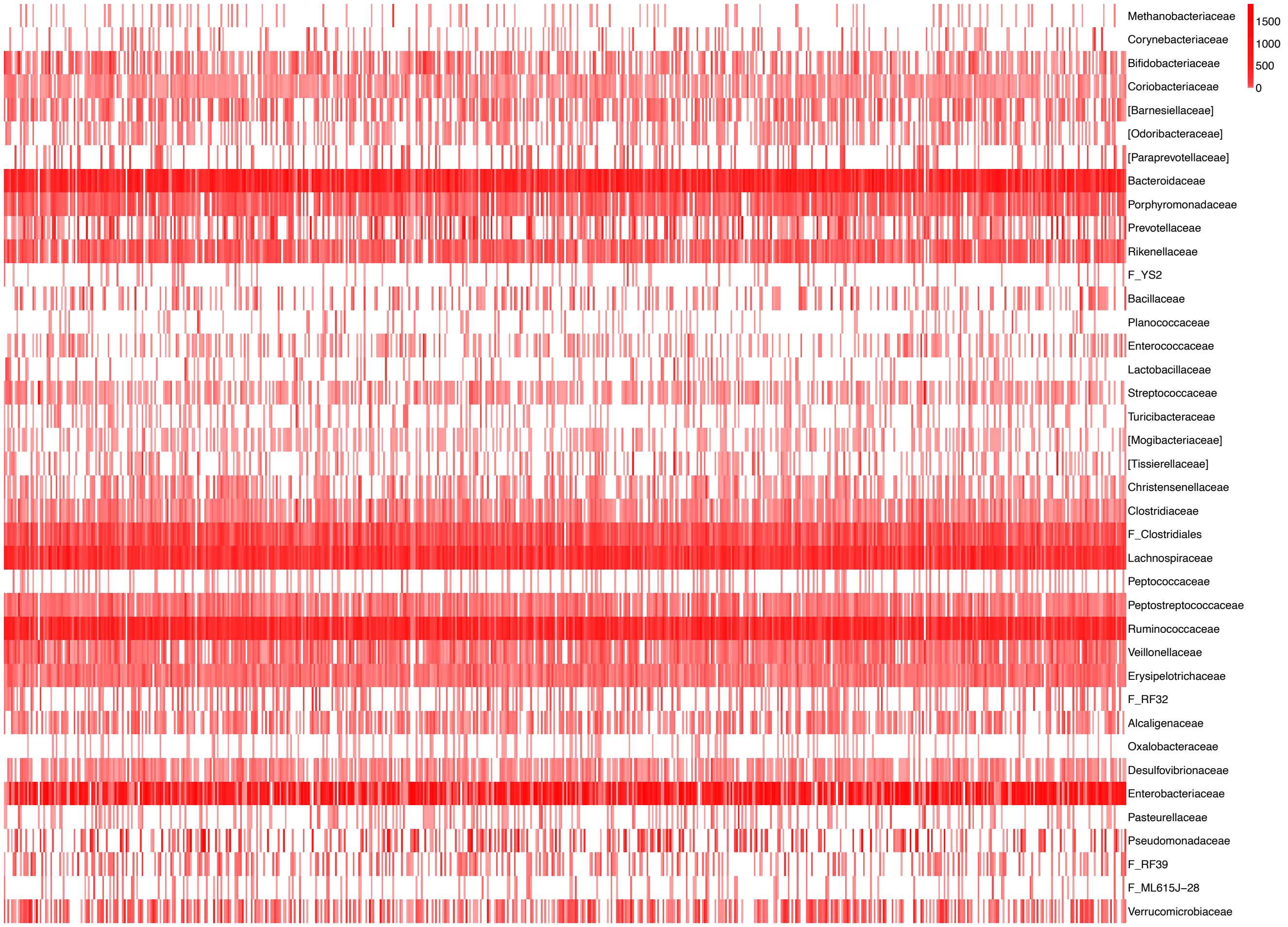


Figure S5: Application – AGP: Heatmap showing the abundance of each of the 39 OTUs at the family level in the taxonomy.

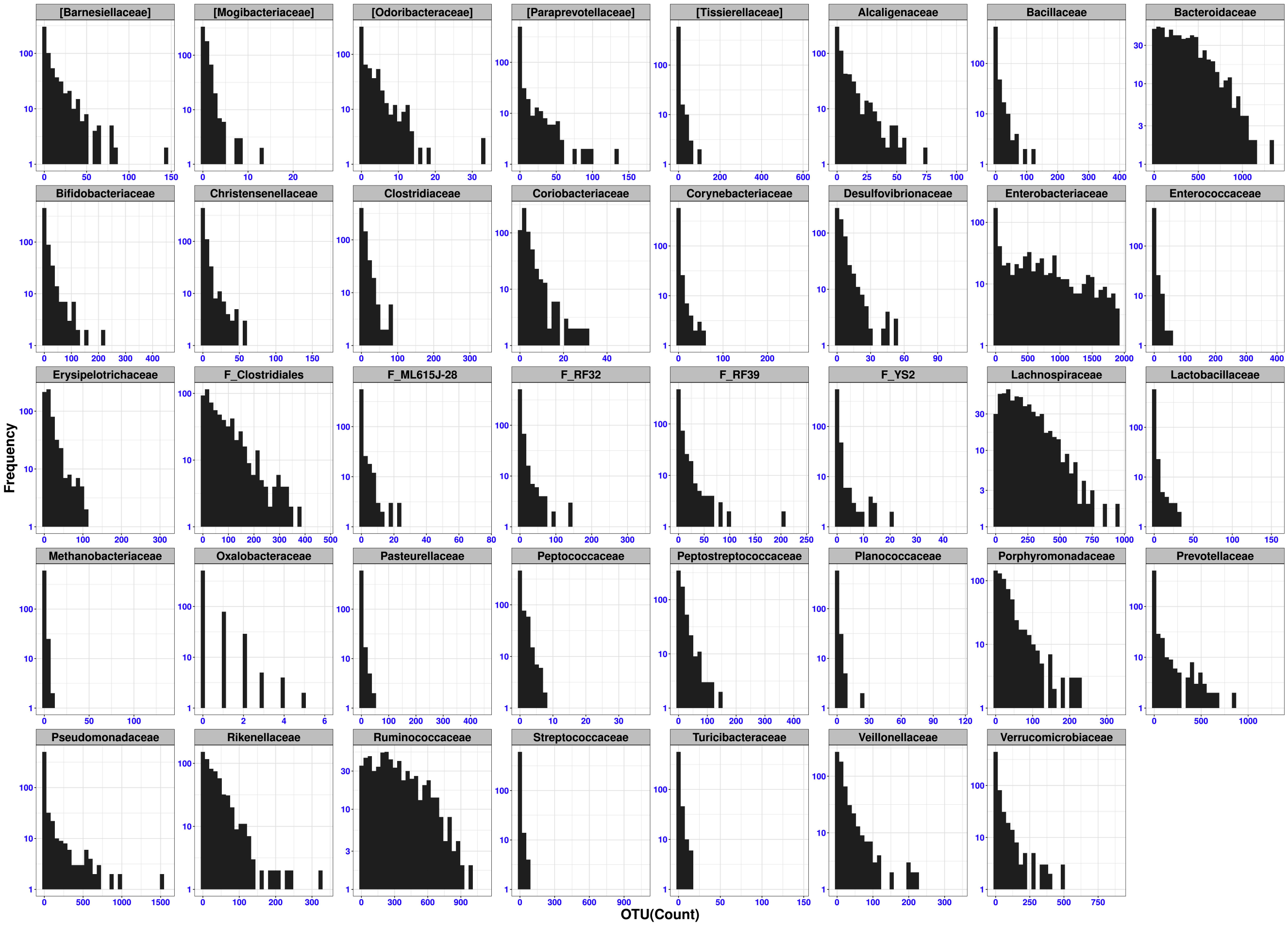


Figure S6: Application – AGP: Histogram showing the distribution of each of the 39 OTUs at the family level in n = 627 samples.

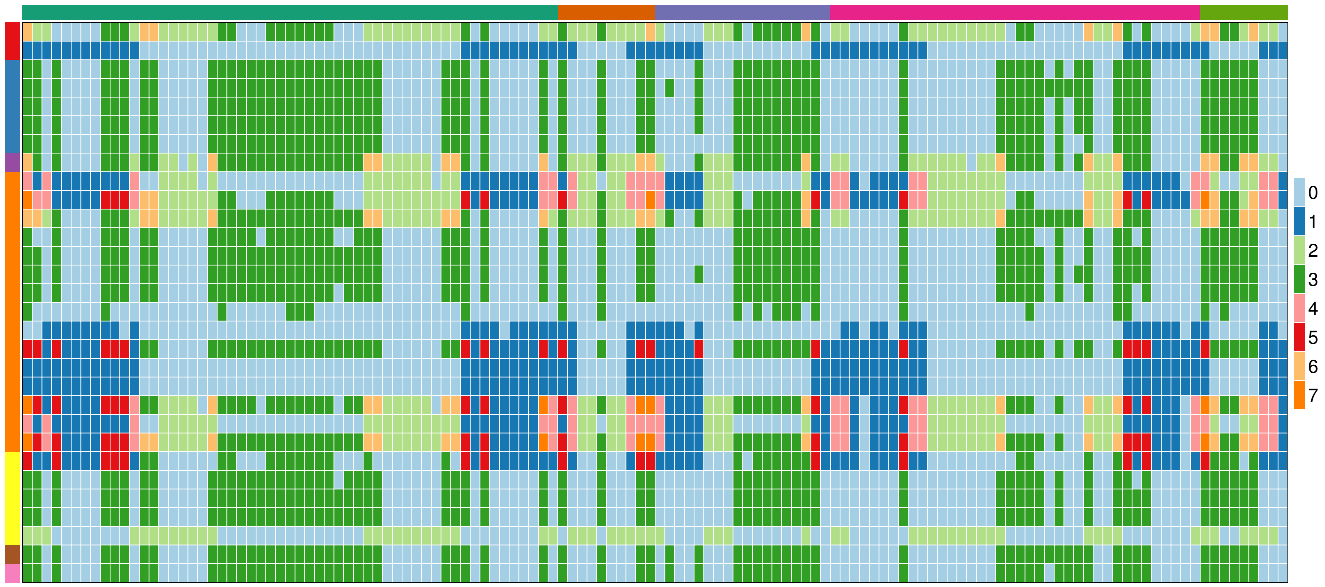


Figure S7: Application – AGP: Color code the contribution of the three unit-rank components in the sparse estimate of the selected rows and columns of the coefficient matrix $\widehat{\mathbf{C}}$. Color code: a) 0 for no contribution, b) 1 for component k =1, c) 2 for component k =2, d) 3 for component k =3, e) 4 for component k ={1,2}, f) 5 for component k ={1,3}, g) 6 for component k ={2,3}, h) 7 for component k ={1,2,3}.

Aditya Mishra, Dipak K Dey, Yong Chen, and Kun Chen. Generalized co-sparse factor regression. *Computational Statistics & Data Analysis*, page 107127, 2020.

R Core Team. *R: A Language and Environment for Statistical Computing*. R Foundation for Statistical Computing, Vienna, Austria, 2019. URL `https://www.R-project.org/`.

Hui Zou and Trevor J. Hastie. Regularization and variable selection via the elastic net. *Journal of the Royal Statistical Society: Series B*, 67(2):301–320, 2005. ISSN 1467-9868. doi: 10.1111/j.1467-9868.2005.00503.x.
